# A ML Framework for Genetic Sequence Identification using 2D Electrical Conductance Probability Distributions from Mixed Data Sets

**DOI:** 10.1101/2024.12.03.626667

**Authors:** Yiren Wang, Hongning Wang, Arindam K. Das, M. P. Anantram

**Affiliations:** Department of Electrical and Computer Engineering, University of Washington, Seattle, WA 98115, USA; Department of Computer Science and Electrical Engineering, Eastern Washington University, Cheney, WA 99004, USA

## Abstract

Genetic sequence identification from electrical characterization of single molecules has emerged as a promising alternative to traditional approaches. Since electrical data on single molecules is extremely noisy due to the limitations of even state-of-the-art approaches, achieving high detection rates is challenging, particularly when the task involves being able to distinguish a sequence from its single base-pair mismatches. To address this issue, we propose an architecture based on combining a convolutional neural network with an ensemble learning method, XGBoost. In addition, four different input feature representations are considered, 1D conductance probability distributions and 2D conductance versus distance probability distributions which can be viewed as images, with or without averaging over the experimental parameters. The with averaging case corresponds to feature matrices derived from mixed datasets. We find that 2D probability distributions are helpful with respect to classifier accuracy, but averaged conductance probability distributions are much more impactful and significantly enhance prediction accuracy. Our quantitative analysis of multiple sequences shows an impressive performance increase of approximately 10% for all sequences. While the basis of our analysis is conductance data of DNA strands for COVID-19 Alpha, Beta, and Delta variants and their single base-pair mismatches, our method is generally applicable to other single-molecule identification based on their conductance.

## Introduction

A DNA sequence is a complex strand of nucleotides that composes the blueprint of every living organism, including viruses. In the past several decades, DNA sequencing techniques have undergone significant development, leading to profound advancements in our understanding of the DNA molecule and its applications in various disciplines, such as molecular biology, genomics, evolutionary biology, virology, medicine, forensics, and so on.^1,2^ Within these disciplines, a key challenge is the accurate identification and classification of genetic sequences and their mutations, particularly those involving single-base pair mismatches.^3^ The ability to accurately identify mutations with single/few base-pair(s) mismatches holds particular significance in the context of viral testing, as evidenced during the recent COVID-19 pandemic.

In recent years, several researchers have explored/proposed single molecule all-electronic methods for DNA/RNA sequence^4–9^ and chemical^10–17^ identification. Broadly speaking, these methods probe the electrical properties of a single molecule by subjecting it to an external voltage source and recording the current (or equivalently, the conductance of the molecule) as the experiment is conducted. Within this paradigm, the identification of sequences rests on the hypothesis that different sequences exhibit different conductance properties, specifically, conductance probability distributions. Such distributions can, therefore, be viewed as “electronic fingerprints” of molecules. Statistical or machine learning methods can then be employed to design classifiers which can produce highly accurate results in near real-time.^18,19^ However, accurate identification of genetic sequences based on their electronic fingerprints can be extremely challenging since even the state-of-the-art methods are inherently stochastic, necessitating multiple experimental passes to collect a “corpus” of data for every molecule.^20–26^ Of course, conductance probability distributions derived from a corpus of data benefit from noise averaging, resulting in more accurate descriptors of the underlying true electronic fingerprints. Nevertheless, from a usability perspective, the holy grail remains highly accurate and robust identification based on data collected from a single experimental pass.

In this paper, we demonstrate that accurate classifiers can be built utilizing the electronic fingerprints of multiple COVID-19 variants and their single base-pair mismatches. The data used in this paper (published in reference^27^) was collected using the single molecule break junction (SMBJ) approach under a range of experimental parameters: applied voltage bias, current amplifier sensitivity, distance between electrodes, and ramp rate (explained in the methods section). As a benchmarking step, we first applied the method we proposed in ^18^ to the data, which yielded low accuracies (83.96%). In this manuscript, we develop a new methodology based on (i) two-dimensional (2D) conductance probability distributions instead of one-dimensional (1D) distributions and (ii) averaged (over the experimental parameters) conductance distributions instead of non-averaged distributions. We trained XGBoost classifiers on 1D distributions and convolutional neural networks paired with XGBoost classifiers on 2D distributions. Our primary research objective was to explore whether any or both of the above variations would better capture the electronic fingerprints of the sequences, which could translate to improved classifier models. Our secondary objectives were: (i) how many experimental passes are needed to construct reliable conductance distributions? and (ii) what role does the value of the applied voltage between the electrodes play, if any? We provide an in-depth analysis of our investigations in the Results section. Our investigations show that: (i) averaged conductance distributions provide an across-the-board enhancement in classifier accuracy (92.09%), (ii) adoption of 2D distributions can be beneficial for some sequences (93.27%) but not all, (iii) 20 experimental passes are adequate to yield classifiers which exceed 90% accuracy for six of the ten sequences considered in this paper, but at least 50 passes are needed for four other sequences, (iv) data collected from experiments conducted at lower voltage biases tend to produce better classifiers, and, (v) higher biases can produce contradictory behavior in classifier accuracy, depending on the sequence type.

## Results and Discussions

The dataset used contains several thousands of current traces corresponding to ten unique 12 base-pair (bp) short DNA segments from one of three variants of the SARS-CoV-2 genome,^27^ as detailed in Table S1 of Supplementary Material. These variants are the Alpha variant B.1.1.7, Beta variant B.1.351, and the Delta variant B.1.167.^28^ Of the ten DNA sequences, three are perfectly matched with the target Alpha, Beta, and Delta variants, referred to as Alpha_PM, Beta_PM, and Delta_PM (perfect match), respectively. The others correspond to sequences of the same length but with a single base pair mismatch, referred to as mismatches (MM). The mismatch sequences denoted by Alpha_MM1, Beta_MM1, and Delta_MM1 are derived from the wild-type SARS-CoV-2 virus, which also exists in some of the unmutated segments of Alpha, Beta, and Delta variants. The mismatch sequences denoted by Alpha_MM2, Beta_MM2, and Delta_MM2 are hypothetical sequences that could produce the same virus protein as the PM sequences.^27^ Table S1 of Supplementary Material shows the exact mutation locations for these sequences. Detailed rationale and conductance data can be found in ^27^.

The current traces are sensitive to the choice of experimental parameters such as the applied voltage bias, ramp rate (i.e., the rate at which the two electrical contacts are separated), and the sensitivity of the current amplifier used to measure conductance. The voltage bias was chosen from the set {0.03, 0.10, 0.15, 0.20} V, the ramp rate applied was from the set {3, 5, 10} V/s, and the current amplifier sensitivity is either 10 nA/V or 1 nA/V (better resolution). As shown in Table S1 of Supplementary Material, this results in a total of 39 experimental datasets of conductance traces for the ten COVID-19 sequences. We will discuss the impact of voltage bias on classifier accuracy in detail later.

We first look for exponentially decaying traces of conductance versus time and discard traces with *R*^2^ < 0.95 as they signify measurements without a molecule in between (invalid data). The threshold of 0.95 was chosen to achieve a balance between filtering out invalid data and retention of adequate data for training and validation (additional details in the Methods section). Then, for each of the ten COVID-19 sequences, we sample *H* current traces randomly and construct a conductance probability distribution (histogram). The detailed procedure for converting a current trace to a conductance trace is provided in the Methods section. While a low value of *H* (ideally one) would be preferable to reduce the experimental burden, conductance histograms constructed from individual or very few current traces are extremely noisy, yielding poor classification accuracy. Our experiments suggest that a reasonable compromise between experimental burden and accuracy is provided by choosing a baseline value of *H* = 30. In this paper, we have experimented with two different sampling methods for the conductance traces, resulting in either a *conductance histogram* or an *average conductance histogram*, and two different histogram representations, 1D or 2D. We discuss these below.

We will first clarify the distinction between a conductance histogram and an average conductance histogram. There are 5 different labels associated with Alpha_MM1: E1, E2, E3, E4, and E5. These labels correspond to different choices of experimental parameters (voltage bias, current amplifier sensitivity, and ramp rate) for the same sequence (see Table S1 of Supplementary Material). When constructing a histogram for Alpha_MM1 from *H* conductance traces, we have two choices: (i) sample all *H* traces from a particular combination of experimental parameters (e.g., E3, corresponding to a bias of 0.1V, a current amplifier sensitivity of 10 nA/V, and a ramp rate of 10 V/s), or (ii) sample *H* traces randomly from all combinations of experimental parameters. A histogram (properly normalized) constructed using the first sampling approach represents a conditional probability distribution of conductance values conditioned on the specific experimental parameters. We refer to such a histogram as a *conductance histogram/probability distribution*. In contrast, a histogram constructed using the second sampling approach represents an average (unconditional) probability distribution, averaged over all combinations of experimental parameters. We refer to such a histogram as an *average conductance histogram/probability distribution*.

Irrespective of the method used to sample *H* conductance traces, we can construct either a 1D histogram or a 2D histogram. We now elaborate on the procedures. As recorded, each conductance trace is a function of time, which can be recalibrated in terms of the distance between the two contacts (as time increases, so does the inter-electrode distance). This conversion procedure is discussed in the Methods section. Since the largest value of the inter-electrode distance is approximately 0.4 nm, we divide the range of distance values, [0, 0.4] nm, into a number of distance bins, which we denote by *N*_bins_Distance_. Similarly, the range of conductance values is divided into a number of conductance bins, which we denote by *N*_bins_. A set of *H* randomly sampled conductance vs. inter-electrode distance traces is then binned along both the distance (*x*) axis and the conductance (*y*) axis, yielding a 2D conductance vs. distance histogram. The 2D histograms, therefore, provide additional information regarding the inter-electrode distance where conductance change occurs, which we have found can be critical in successfully distinguishing between certain SARS-CoV-2 variants. By summing over the distance axis, we can marginalize the 2D histogram, yielding a 1D conductance histogram. Please refer to Figures S1, S2, S3 and S4, S5, S6 of Supplementary Material for 1D and 2D histograms, respectively, for each experimental dataset. Conceptually, the 2D histograms can be viewed as equivalent to a time-frequency analysis (e.g., short-time Fourier Transform), whereas 1D histograms can be viewed as equivalent to a Fourier Transform. We wish to emphasize that this analogy is merely pedagogical, and we do not convert the data to the frequency domain.

Next, we construct 1000 histograms for each sequence, 70% of which are used for classifier training and validation, while the rest are used for testing. We present results from four different input data representations, which are dictated by (i) choice of 1D histograms or 2D histograms, and (ii) choice of conductance histograms or average conductance histograms. Our main motivation for studying the performance of 2D distributions is to examine whether such distributions are any more informative than 1D distributions. Table 1 summarizes the four possible input data representations, the ML algorithm used, and the average (over 100 trials) accuracy for a *baseline classifier*. We define a baseline classifier as one which is characterized by the following parameters: (i) prefiltering with *R*^2^ ≤ *β* = 0.95, (ii) histograms from *H* = 30 current traces, and (iii) number of conductance bins *N*_*bins*_ = 600 (for both 1D and 2D histograms) and number of distance bins *N*_bins_Distance_ = 10 for 2D histograms (equal to 1 for a 1D histogram).

**Table 1:** Summary of four different input data representations, ML algorithms used, and corresponding classification accuracies.

| Approach | Histogram dimensionality | Average conductance histogram? | ML algorithm | Average accuracy |  |
| --- | --- | --- | --- | --- | --- |
| | | | | $H = 30$<br>(baseline) | $H = 10$ |
| A1 <sup>18</sup> | 1D | No | XGBoost | 83.96% | 74.50% |
| A2 | 1D | Yes | XGBoost | 92.09% | 82.60% |
| A3 | 2D | No | CNN + XGBoost | 85.60% | 75.96% |
| A4 | 2D | Yes | CNN + XGBoost | 93.27% | 85.57% |

Average accuracy over 100 trials for sample sizes *H* = 10 and 30 are also included in Table 1. In general, the performance of the classifier depends on the choice of the parameters *β, H*, and *N*_bins_, of which the sample size parameter *H* has the most pronounced effect.

For 1D histograms, we have used the XGboost classifier algorithm. Two critical hyperparameters within an XGboost framework are the number of trees/estimators, *N*_*est*_, and the depth of each tree/estimator, *D*_*est*_. Based on extensive hyperparameter optimization, we chose *N*_*est*_ = 200 and *D*_*est*_ = 2. For 2D histograms, we use a Convolutional Neural Network (CNN) for feature extraction. The output from the final layer of the CNN is fed to an XGBoost classifier, as in the 1D case. We also experimented with a traditional CNN paired with a fully connected layer classifier, but its results are not as accurate as a CNN + XGBoost approach. For example, the CNN-only approach shows more than a 5% decrease in classification accuracy on the Alpha variants compared to a CNN + XGBoost approach (see Figure S7 of Supplementary Material). Additional details on the XGBoost and CNN classifiers are available in the Methods section.

### Performance of baseline classifier

In this section, we discuss the performance of a *baseline classifier* by comparing in detail the differences between approaches A1 (worst performing) and A4 (best performing) indicated in Table 1. We observe that for small sample sizes such as *H* = 10, approach A4, which utilizes 2D average conductance histograms and a CNN + XGBoost classifier, always has a similar or slightly better overall accuracy than approach A2, which utilizes 1D average histograms and an XGBoost classifier. We also observe that the accuracy improves by 7-8% when average conductance histograms (whether 1D or 2D) are used, compared to conductance histograms which are conditioned on experimental parameters. This can be verified by comparing the accuracies of approaches A1 vs. A2 and approaches A3 vs. A4, shown in Table 1.

Our primary research objective is as follows: “Can the PM (perfect match) and MM (mismatch) sequences for each COVID-19 variant (Alpha, Beta, Delta) be identified with high accuracy?”. For each of the three variants, we treat the PM and MM sequences as different target classes. This results in three classifiers, one each for the three variants; specifically, (i) 3-class classifiers for the Alpha and Delta variants since these variants have one PM class and two MM classes, and (ii) a 4-class classifier for the Beta variant since this variant has one PM class and three MM classes.

Figure 1 shows the confusion matrices for the Alpha, Beta, and Delta baseline classifiers; panels (a,b,c) correspond to Approach A1,^18^ while panels (d,e,f) correspond to Approach A4. Please refer to Figure S8 of Supplementary Material for the confusion matrices of all four approaches. Comparing Figure 1(a) and 1(d), we see that 1D histograms work reasonably well (≈ 90% accuracy) for distinguishing the Alpha_MM2 class, but there is major confusion between the Alpha_MM1 and Alpha_PM classes (the 1D histograms for these classes are very similar). However, usage of 2D average histograms provides an across-the-board improvement in classification accuracy, approximately 8% for Alpha_MM2 and 13-15% for the Alpha_MM1 and Alpha_PM classes. The case for the Beta classifier is rather interesting. Comparing Figure 1(b) and 1(e), we can see that although 1D histograms can distinguish the Beta_PM class perfectly, the accuracy is approximately 75-78% for the three mismatch classes. This suggests that 1D histograms for the Beta_PM class must be strikingly dissimilar to the three mismatch classes, which is indeed the case, as evidenced by Figure S2 of Supplementary Material. Usage of 2D average histograms boosts the accuracies of the mismatch class Beta_MM2 by about 20% and Beta_MM3 by about 12%, although there is still room for improvement as far as Beta_MM1 and Beta_MM3 classes are concerned (roughly 15% confusion between these two classes). Finally, comparing Figure 1(c) and 1(f), we observe that 2D average histograms provide an average accuracy boost of approximately 4-10% for the Delta classes.

**Figure 1.**
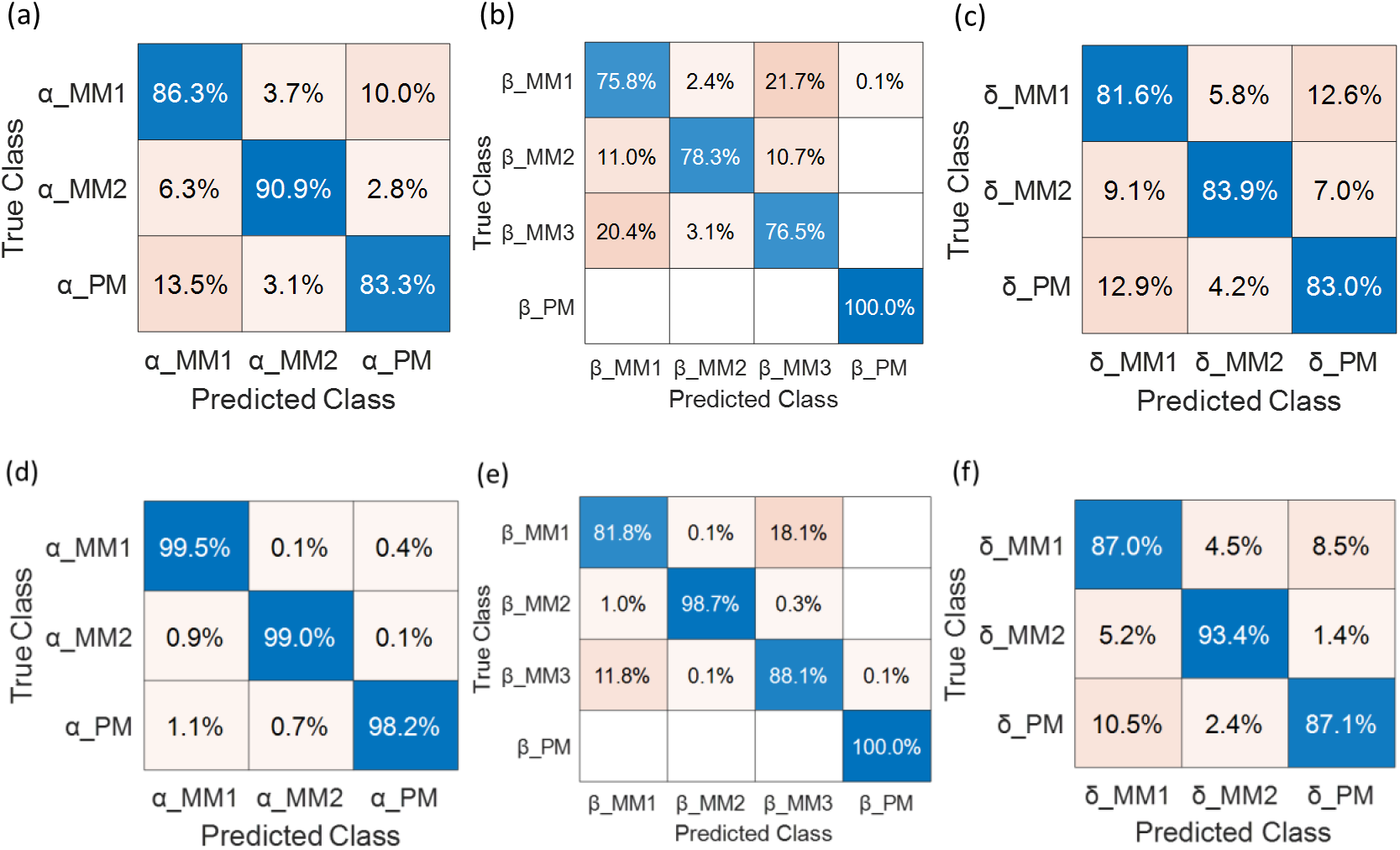
Confusion matrices for baseline classifiers: (a,b,c) for approach A1 (d,e,f) for approach A4. The differences between these two approaches are summarized in Table 1.

### Performance analysis of baseline classifiers w.r.t sample size *H*

In this section, we address the impact of the parameter *H*, which tends to affect classifier accuracy the most. Intuitively, we expect larger values of *H* to yield better classifier models since the relatively high stochasticity inherent in the current traces will tend to be ‘averaged out’ as *H* increases, revealing the underlying true probability distribution. From a ‘usability’ perspective, a value of *H* = 1 is ideal since it implies that a sequence determination can be made at run time after every experiment. However, the state-of-the-art experimentation currently precludes operating at an ideal value of *H*.

Figure 2 shows the overall classification accuracy for all four approaches indicated in Table 1 with respect to the sample size, *H*, while keeping *β* = 0.95, *N*_*bins*_ = 600, and *N*_bins_Distance_ = 10. A similar plot over an extended range of *H, H* ∈ [10,100], appears in Figure S9 of Supplementary Material. The reason why we study all four approaches in this section is that Figure 2 provides some clues regarding the relative efficacies of the two data representational attributes, (i) 2D vs. 1D histograms and (ii) average histogram or not, as discussed in the subsequent paragraphs.

**Figure 2.**
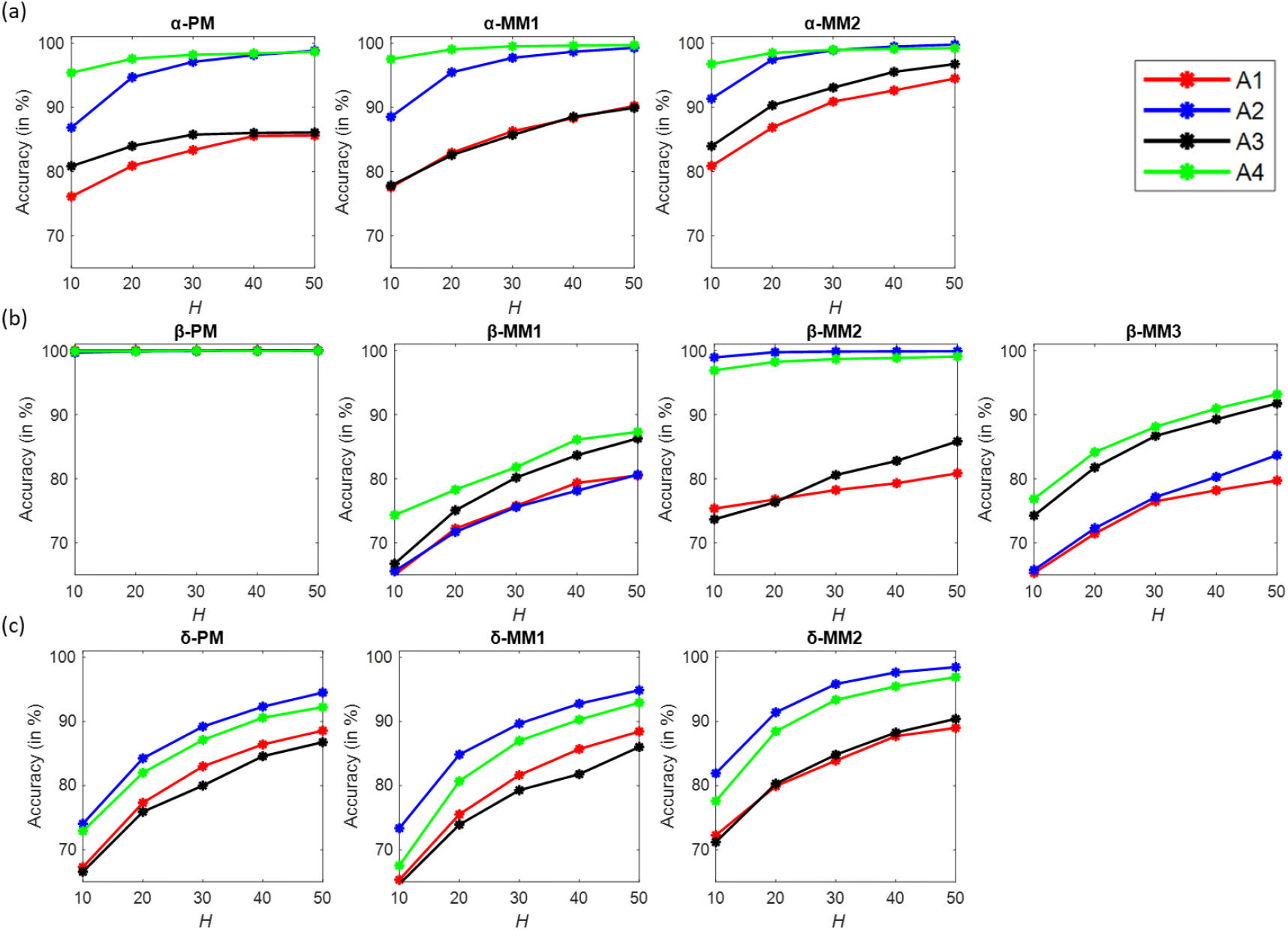
Performance analysis of baseline classifiers with respect to H for the four approaches indicated in Table 1. (a) Alpha sequences, (b) Beta sequences, and (c) Delta sequences. The same color scheme has been used to distinguish between the four approaches.

From the top row in Figure 2, we observe that for the Alpha variant, approach A4 is better than A2, which is significantly better (about 10%) than A3 and A1. This suggests that averaged histograms are more impactful in the case of Alpha variants, particularly for the Alpha_PM and Alpha_MM1 sequences. We also observe that approach A4 is approximately 95% accurate for Alpha variants when *H* = 10, compared to ≈ 87% for approach A2. However, as *H* increases, the two approaches become comparable and are visually indistinguishable for *H* ≥ 40. Therefore, for the Alpha variant, we conclude that approach A4 is preferable over approach A2, particularly for smaller values of *H*.

From the bottom row in Figure 2, we observe that the plots for the Delta variant are slightly different from the Alpha variant. In this case, approach A2 is the best overall, followed by approach A4, even for smaller values of *H*. In fact, for the Delta_MM1 sequence, A2 is approximately 6% better than A4 when *H* = 10. We also observe that neither of the two approaches can provide a 95% classification accuracy or better (our intended benchmark) for all three Delta variants even when *H* = 50. In general, we have found that Delta variants are harder to classify than Alpha variants.

Finally, the case for the Beta variant turns out to be significantly different than the Alpha and Delta variants. From the middle row in Figure 2, we observe that the Beta_PM class is easiest to classify, reaching almost 100% accuracy at *H* = 10, irrespective of which of the four approaches is used. For the Beta_MM1 and Beta_MM3 sequences, averaged 2D histograms (approach A4) perform marginally better than non-averaged 2D histograms (approach A3), and either of these two approaches is better than approaches A1 and A2 which utilize 1D histograms. This phenomenon is similar to what we observed for the Alpha sequences. For Beta_MM2, however, approach A2 performs marginally better than approach A4, as is the case for the Delta variant sequences. In fact, if we compare the blue/green and red/black curves for Beta_MM2, we observe that averaged histograms provide almost 23-24% improvement over non-averaged histograms when *H* = 10.

We now summarize the observations/inferences from the preceding discussion. While we embarked on this analysis with the goal of being able to state unequivocally which of the four approaches is best, it turns out that no straightforward answer exists. Disregarding the Beta_PM sequence for which all four approaches provide 100% accuracy even for *H* = 10, we find that averaged 2D histograms (approach A4) perform best for all three Alpha sequences and the Beta_MM1/MM3 sequences. However, averaged 1D histograms (approach A2) perform best for all three Delta sequences and the Beta_MM2 sequence. Our investigations suggest that while averaged conductance histograms are preferred over conditional histograms when experiments on COVID-19 variants and their mismatches are conducted using a range of experimental parameter values, the choice of a 1D or 2D histogram is sequence-dependent. The exact reason why some sequences benefit from a time/tip distance indexed conductance distribution, but not all, is not understood at this point and is subject to further theoretical investigation.

### Impact of applied bias on classifier accuracy

So far, we have discussed the performance of our baseline classifiers and their sensitivities to the *histogram related parameter, H*. In this section, we explore how the *SMBJ experimental parameter*, voltage bias, impacts classification accuracy. Obviously, this analysis necessitates a fine-grained data labeling scheme that is cognizant of the sequence and the voltage bias. Since the majority of datasets in Table S1 of Supplementary Material have been recorded using a current amplifier with a sensitivity of 10 nA/V and a ramp rate of 10 V/s, we first filtered Table S1 of Supplementary Material based on these two parameters. Subsequently, we combined datasets which pertain to the same sequence and applied bias.

For example, for the Alpha variant, the filtered datasets are E3, E4, E5, E8, E9, E14, and E15, all of which were recorded using a 10 nA/V amplifier and a 10 V/s ramp rate, although they differ in the applied bias (0.10 V, 0.15 V, and 0.20 V). We refer to this subset of seven datasets as the Alpha variant for our analysis in this section. For the Beta variant, the filtered datasets are E16, E17, E18, E21, E22, E23, E24, and E27, which differ in the applied bias (0.05 V and 0.10 V). However, since E16 and E18 correspond to the same sequence (Beta_MM1) and a bias value of 0.10 V, we combined these two datasets. Similarly, E23 and E24 were combined since they both correspond to Beta_MM3 and a bias value of 0.10 V. This resulted in six datasets for the Beta variant, E16 ∪ E18, E17, E21, E22, E23 ∪ E24, and E27, where ∪ denotes the set union. Following an identical procedure, we end up with eight datasets for the Delta variant, E51 ∪ E52, E53 ∪ E54, E55, E56 ∪ E57, E58, E59 ∪ E60, E61, and E62.

Next, we generated histograms using approach A3 (2D non-averaged histograms, see Table 1) and *H* = 100. A reasonably large value of *H* was chosen to minimize the effect of sampling-induced uncertainties on classifier performance. It is important to note that each such histogram represents a *unique* combination of the tuple (sequence, voltage bias). These histograms described in the previous paragraph are then used to train and test independent classifiers for each of the three variants (Alpha, Beta, and Delta): (i) a 7-class classifier for the Alpha variant, (ii) a 6-class classifier for the Beta variant, and (iii) an 8-class classifier for the Delta variant. Table 2 shows the classifier accuracies for Alpha, Beta, and Delta variants as a function of applied bias. The confusion matrices for these three classifiers are shown in Figures S10, S11, and S12 of Supplementary Material. Results for classifiers trained on histograms with *H* = 30 are shown in Table S2 of Supplementary Material.

**Table 2:**
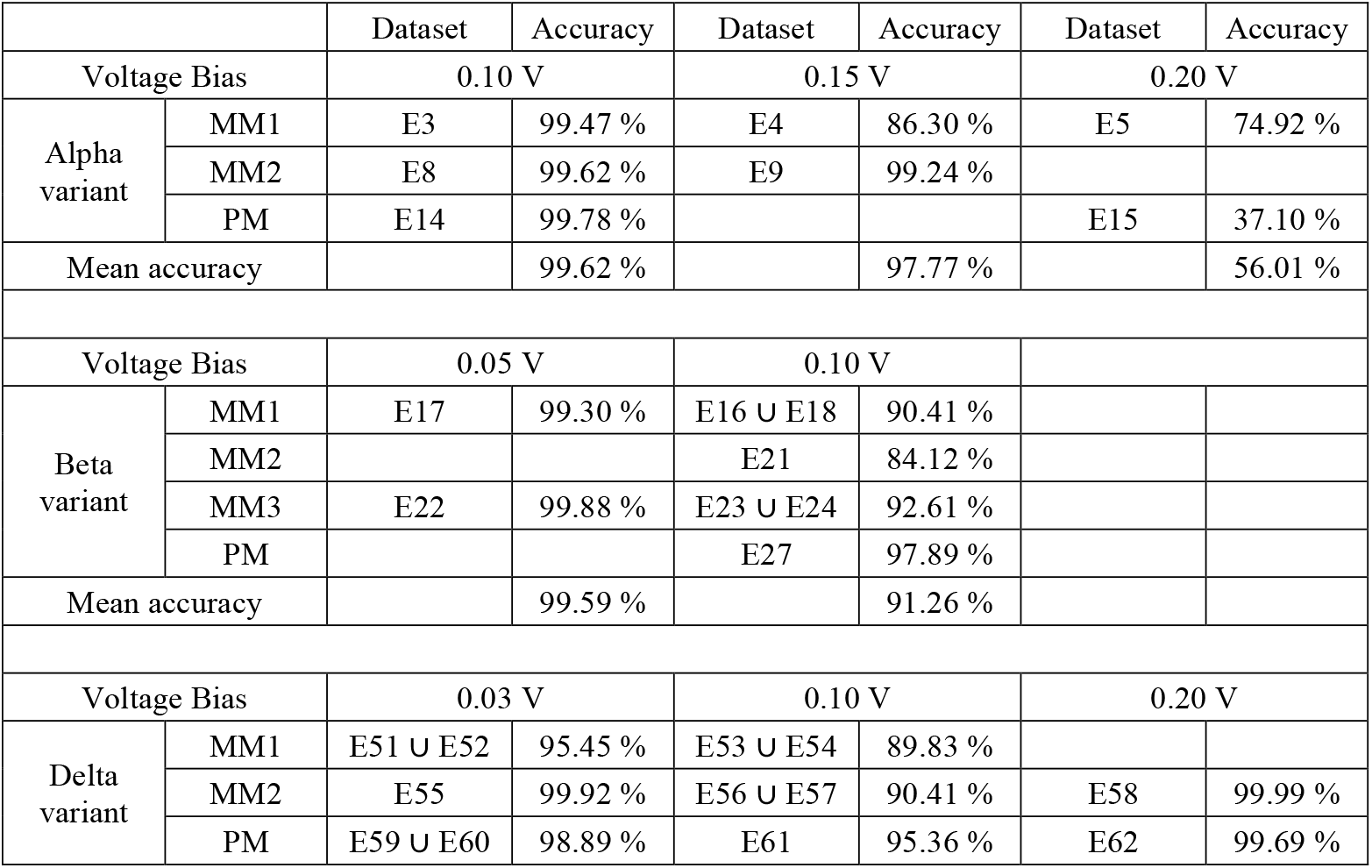
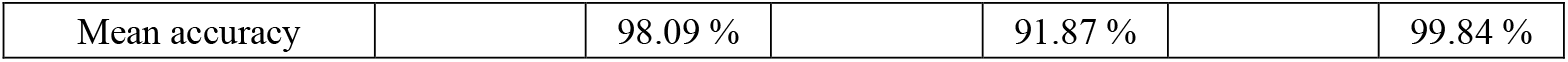
Classifier accuracies for Alpha, Beta, and Delta variants as a function of applied bias (histograms generated using H = 100). All datasets referenced in this table were recorded using a current amplifier of sensitivity 10 nA/V and a ramp rate of 10 V/s.

A general observation from Table 2 is that lower biases result in higher accuracies if we restrict ourselves to a moderate bias range [0.03, 0.15] V. This is true for all three variants. However, when the bias is 0.20 V, we observe contradictory behaviors for the Alpha and Delta variants. While the accuracy drops precipitously for the Alpha sequences E5 (Alpha_MM1) and E15 (Alpha_PM) at a bias of 0.20 V, it approaches almost 100% for the Delta variant sequences E58 (Delta_MM2) and E62 (Delta_PM). This contradictory behavior can be validated by using an information-theoretic criterion, namely the Jensen-Shannon metric, as explained in Section 5 of Supplementary material (see Figure S13 in particular).

The preceding discussion regarding the contradictory behavior exhibited by the Alpha and Delta variant sequences at a bias of 0.20 V raises an interesting question: are some DNA sequences naturally more susceptible to higher applied bias? Could it be due to some structural deformations within the sequence, which are induced by higher bias values? We leave this as an open question for now, pending additional theoretical investigation.

## Conclusions

The identification of genetic material and chemical detection from the conductance of single molecules has been pursued for over a decade. The noisy nature of the experimental measurements on single molecules has been a challenge in enabling robust identification. Further, physics-based modeling has failed to theoretically discern between molecules by accounting for the possible contact configurations and environmental fluctuations. In this paper, we propose a methodology based on multiple machine learning techniques and two different methods for data representation to identify DNA strands with high accuracy. Accompanying the developed method is the release of a software package that can be used in a wide range of molecular conductance data.

Experimentally, the distance between the metal electrodes is varied during an SMBJ run, and conductance is measured as a function of time. As a result of the intrinsically noisy nature of the experiments, the conductance values are not completely reproducible if experiments are repeated on the same molecule. This prompted us to represent the data as conventional 1D conductance probability distributions and, additionally, as a 2D probability distribution, which depends on both the conductance value and inter-electrode separation. We studied the accuracy of detection both with and without averaging the conductance distribution over experimental parameters. In conjunction with 2D distributions, we have experimented with a convolutional neural network paired with an XGBoost classifier. Our investigations reveal that: (i) averaged conductance distributions have a significant impact on classifier accuracy, boosting it by as much as 21%, (ii) 2D conductance distributions are beneficial for some sequences, but not all, and (iii) the backend XGBoost classifier whose input is data from a convolutional neural network provides better accuracy than a densely connected classifier usually adopted with a convolutional neural network. Finally, the classifier models were used to study the impact of the applied voltage between the two electrodes on accurate sequence detection. We find prima facie evidence that relatively large biases can be detrimental to the development of accurate classifier models. Overall, our classification approach exhibits significant potential for accurate, amplification-free DNA sequence identification. Importantly, while our analysis focused on COVID-19 data, the computational method proposed in this paper should be of use in analyzing arbitrary single-molecule conductance data for sequence identification.

## Methods

A series of re-processing steps were carried out to convert the experimentally obtained ‘raw’ current traces to suitable conductance traces, which are then used to derive the conductance histograms for training our ML models. Figure 3 summarizes the sequence of pre-processing steps. Figure 4(a) shows a DNA between the two electrodes, where the applied bias drives the current flow, the ramp rate determines how fast the distance between the electrodes increases, and the preamplifier can also be varied in the experiments. The recorded current values are the outputs of a preamplifier, which depends on the selection of a current amplifier of either 1 nA/V or 10 nA/V sensitivity.^27^ Since an amplifier of 1 nA/V sensitivity has a noise floor of 1 *p*A and an upper limit of 10 *n*A, we first clipped all current traces recorded with such an amplifier to the range [1 *p*A − 10 *n*A]. Since a 10 nA/V amplifier has a noise floor of 10 *p*A and an upper limit of 100 *n*A, we clipped all current traces recorded with such an amplifier to the range [10 *p*A − 100 *n*A].

**Figure 3.**
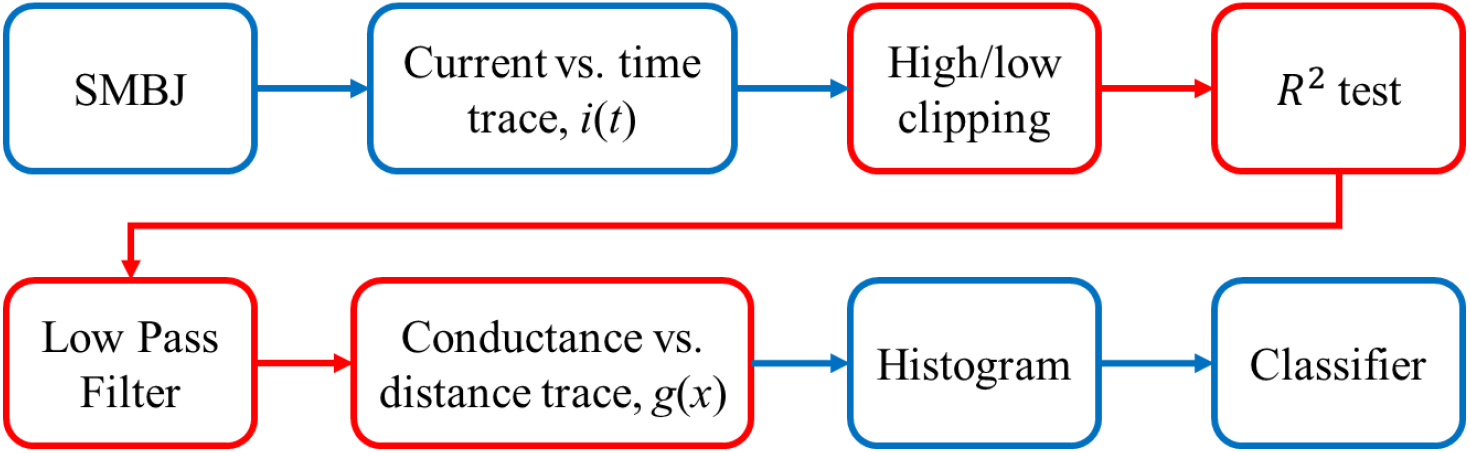
Sequence of pre-processing steps (shown within red boxes) for converting raw current traces to conductance traces.

**Figure 4.**
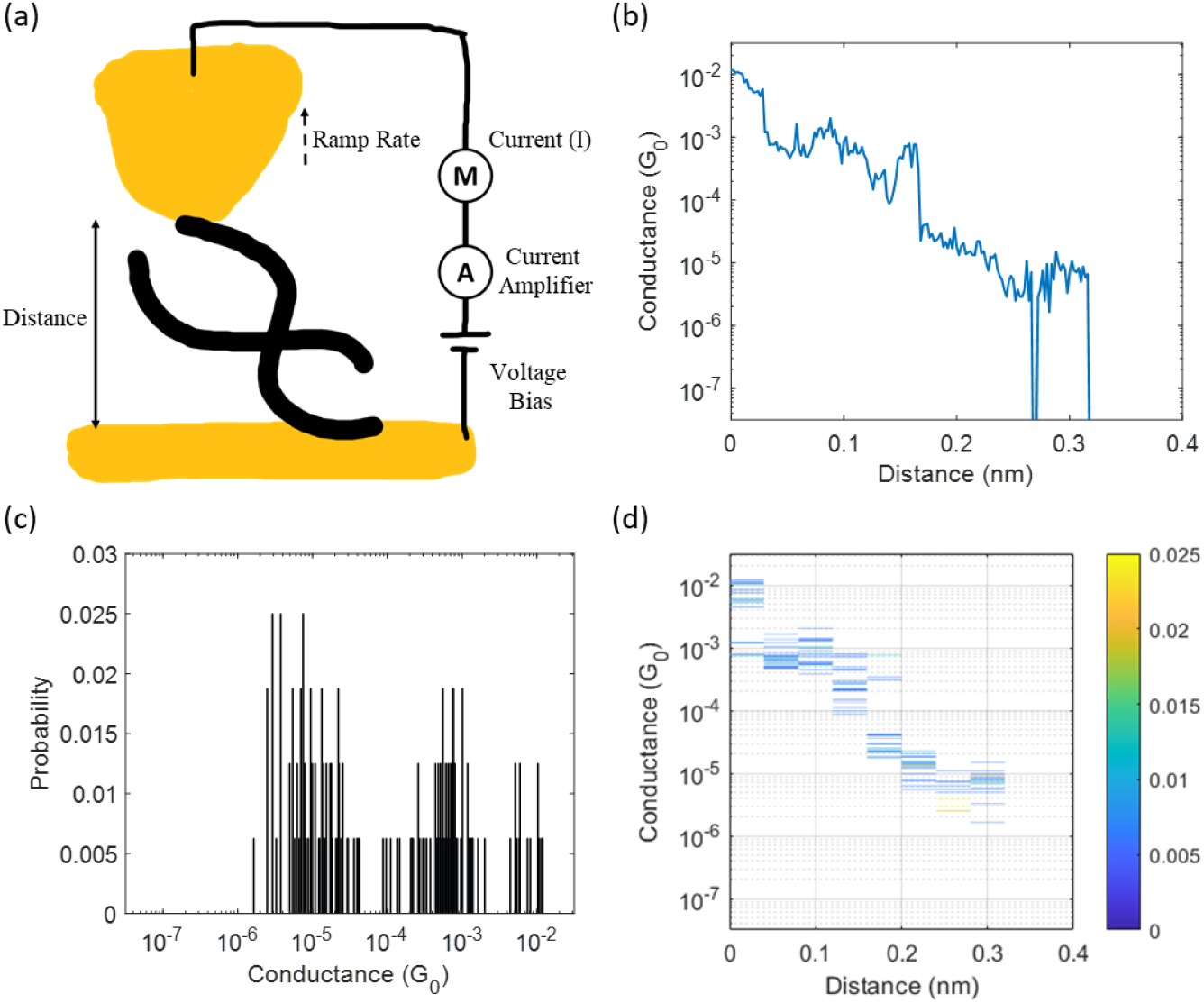
(a) Schematic of the SMBJ experimental method, (b) a single conductance versus distance trace, (c) derived 1D conductance histogram, and (d) derived 2D conductance histogram.

Next, we fit an exponential regression model, *y* = *a* · *e*^−*b*·*x*^, *b* > 0, to each high/low clipped current trace and used the *R*^2^ statistic from the fit to accept/reject the trace. Specifically, if the computed *R*^2^ statistic is greater than some chosen threshold, *β*, we reject the trace as ‘invalid’ (experiment conducted with no molecular binding); otherwise, the trace is accepted. Even though the standard error of regression is a better choice for nonlinear regression models, we determined through extensive experimentation that the *R*^2^ statistic works well for our purposes. We started with about 2000−5000 traces for each of the 39 datasets. For the Alpha variant, we found that approximately 60−85% of the data were accepted at *β* = 0.95. There were two exceptions: E7 corresponding to Alpha_MM2 (42% accepted) and E13 corresponding to Alpha_PM (20% accepted). For the Beta variant, other than E25, E26, and E27 (corresponding to Beta_PM), 56-86% of the current traces were accepted at *β* = 0.95. For Beta_PM, the acceptance rate was 20-23%. For the Delta variant, we found that approximately 71−90% of the data were accepted at *β* = 0.95.

In the third step, all accepted current traces were low-pass filtered to mitigate high-frequency noise. Since the sampling frequency for all 39 datasets is 10kHz, we used a low pass filter with a cutoff frequency of 3 kHz.

In the fourth step, all current traces were converted to conductance traces using the equation below:

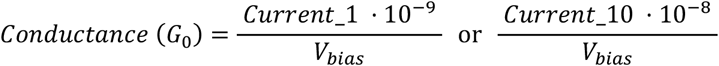

where Current_1 is in units of 1 nA, Current_10 is in units of 10 nA, and *V*_*bias*_ is the bias voltage in volts. The unit *G*_0_ in the above equation is the *conductance quantum* and is defined as follows: 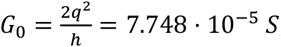, where *q* is the elementary charge and *h* is Planck’s constant. For our datasets, the conductance values turn out to be in the range [10^−7.5^ − 10^−1.5^] *G*_0_.

The conductance versus time (*g*(*t*)) traces obtained in the previous step are recalibrated as conductance versus distance (*g*(*x*)) traces using the eq. below:

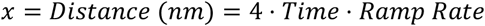

where each time step is 10^−4^ s corresponding to the 10kHz sampling frequency. Ramp rate, in V/s, is the voltage applied to control the rate at which the distance between the two electrodes changes. The distance between the electrodes is in units of nm, and most measurements have a distance value of less than 0.4 nm. Therefore, we choose [0 – 0.4] nm as the range of distance values for creating histograms.

Finally, the conductance vs. distance traces is sampled to compute 2D and 1D histograms (viewed as probability distributions), which are used to train/test the classifiers. Figure 4 shows sample 2D and 1D histograms constructed from a single conductance trace along with the figure of the experimental set up and the parameters.

### Machine Learning Models

XGboost^29,30^ is a fast and scalable implementation of a gradient boosted decision tree framework.^31^ Gradient boosting is an ensemble learning method wherein weak base learners (usually decision trees) are added sequentially, one at each iteration, to minimize a suitably defined loss function evaluated on the previous learner. Within XGboost, the loss function is cross entropy for multiclass classification problems. We used the Python implementation of the XGboost package.^32^ Two critical hyperparameters within an XGboost framework are the number of trees/estimators, *N*_*est*_, and the depth of each tree/estimator, *D*_*est*_.

Optimal values of these parameters were determined based on an exhaustive grid search on values indicated below:

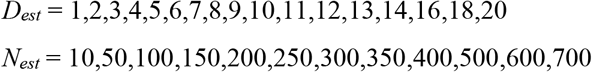

Based on validation accuracies, we determined that reasonable values are (*N*_*est*_ = 200, *D*_*est*_ = 2) or (*N*_*est*_ = 300, *D*_*est*_ = 3). We chose the first parameter combination for all simulations to reduce the computational time. Other hyperparameters were explored to optimize the training time while maintaining a high validation accuracy.

Convolutional Neural Network (CNN) is a deep learning method designed for image and spatial data analysis.^33^ CNNs leverage processes like those in animal sight to detect high and low-level information from image data to make inferences. A CNN consists of multiple building blocks, including convolutional, pooling, and batch normalization layers, to detect features in the input data. The back end of a CNN is typically a fully connected layer (or layers) which classify the extracted features into target classes.

We used the Tensorflow implementation of CNN, with a learning rate based on the Adam optimizer.^34^ We employ categorical cross entropy for the multiclass classification. Figure 5 shows the architecture of our stacked CNN feature extractor and XGBoost model. The inputs to the XGBoost model are the outputs of the flattening layer from a CNN model trained with a fully connected classifier. As mentioned in the Results section, adopting an ensemble classifier model (XGBoost) instead of a traditional fully connected classifier results in an enhancement of more than 5% classifier accuracy. We reckon this is due to the inherent stochasticity of the experimental data, which introduces a variance in the conductance probability distributions. Ensemble learning methods, in general, are better suited to data which are noisy and exhibit high variance.

**Figure 5.**
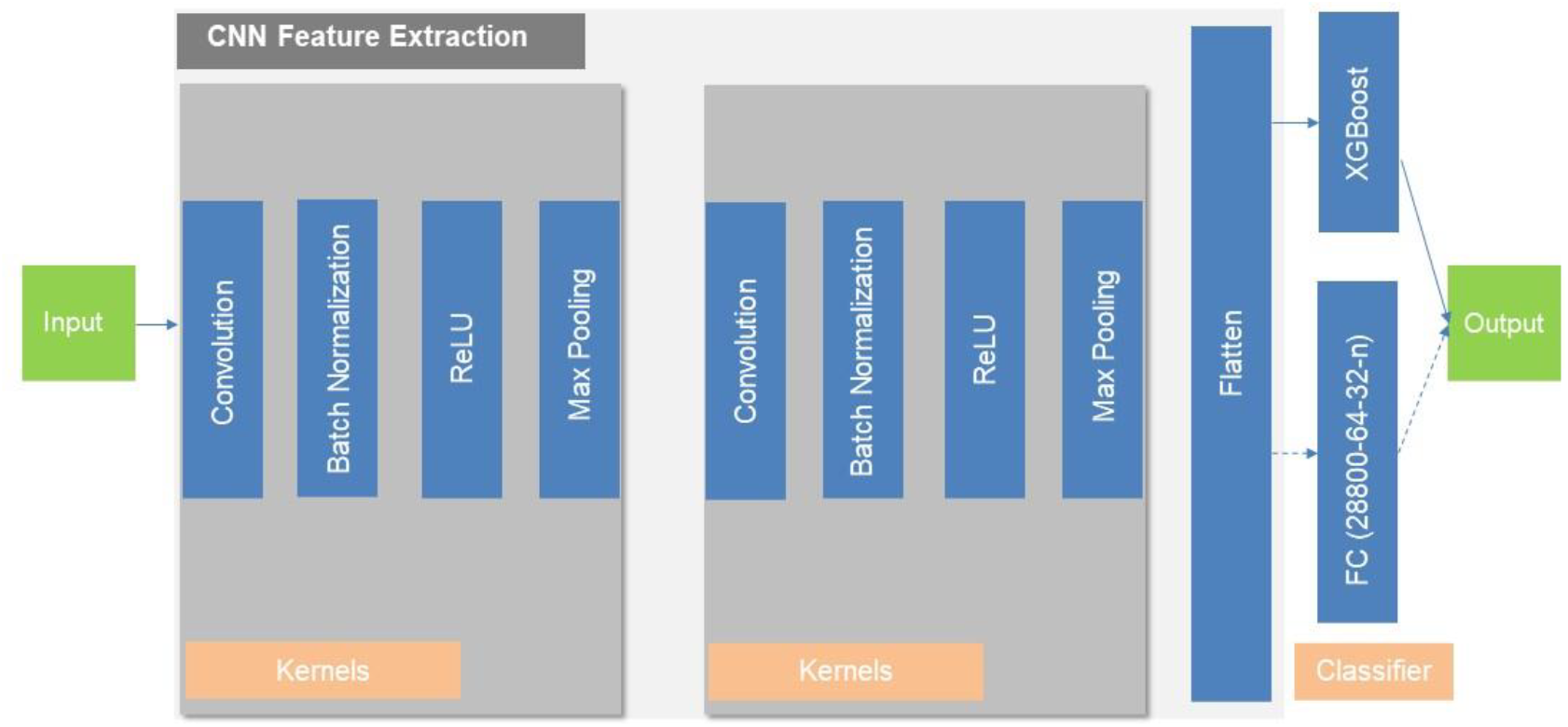
Sequence of components in the stacked CNN and XGBoost model. FC stands for fully connected layer.

All codes/data used for generating the results in this paper are available as a Python package.^35^

## Acknowledgment

We acknowledge NSF grant numbers 2317693 (FMSG) and 2027165 (SemiSynBio).

## References

(1) Ritchie, J. Probabilistic DNA Evidence: The Laypersons Interpretation. Australian Journal of Forensic Sciences 2015, 47 (4), 440–449. 10.1080/00450618.2014.992472.

(2) Dewey, F. E.; Pan, S.; Wheeler, M. T.; Quake, S. R.; Ashley, E. A. DNA Sequencing Clinical Applications of New DNA Sequencing Technologies. Circulation 2012, 125 (7), 931–944. 10.1161/CIRCULATIONAHA.110.972828.

(3) Vogelstein, B.; Papadopoulos, N.; Velculescu, V. E.; Zhou, S.; Diaz, L. A.; Kinzler, K. W. Cancer Genome Landscapes. Science (1979) 2013, 339 (6127), 1546–1558. 10.1126/science.1235122.

(4) Li, Y.; Artés, J. M.; Demir, B.; Gokce, S.; Mohammad, H. M.; Alangari, M.; Anantram, M. P.; Oren, E. E.; Hihath, J. Detection and Identification of Genetic Material via Single-Molecule Conductance. Nat Nanotechnol 2018, 13 (12), 1167–1173. 10.1038/s41565-018-0285-x.

(5) Pattiya Arachchillage, K. G. G.; Chandra, S.; Piso, A.; Qattan, T.; Artes Vivancos, J. M. RNA BioMolecular Electronics: Towards New Tools for Biophysics and Biomedicine. J Mater Chem B 2021, 9 (35), 6994–7006. 10.1039/D1TB01141C.

(6) Xu; Zhang; Li; Tao. Direct Conductance Measurement of Single DNA Molecules in Aqueous Solution. Nano Lett 2004, 4 (6), 1105–1108. 10.1021/nl0494295.

(7) Nishino, T.; Bui, P. T. Direct Electrical Single-Molecule Detection of DNA through Electron Transfer Induced by Hybridization. Chemical Communications 2013, 49 (33), 3437. 10.1039/c3cc38992h.

(8) Gunasinghe Pattiya Arachchillage, K.G.; Chandra, S.; Williams, A.; Rangan, S.; Piscitelli, P.; Florence, L.; Ghosal Gupta, S.; Artes Vivancos, J. M. A Single-Molecule RNA Electrical Biosensor for COVID-19. Biosens Bioelectron 2023, 239, 115624. 10.1016/j.bios.2023.115624.

(9) Dief, E. M.; Darwish, N. SARS-CoV-2 Spike Proteins React with Au and Si, Are Electrically Conductive and Denature at 3 × 10 ^8 V m -1^ : A Surface Bonding and a Single-Protein Circuit Study. Chem Sci 2023, 14 (13), 3428–3440. 10.1039/D2SC06492H.

(10) Dief, E. M.; Low, P. J.; Díez-Pérez, I.; Darwish, N. Advances in Single-Molecule Junctions as Tools for Chemical and Biochemical Analysis. Nat Chem 2023, 15 (5), 600–614. 10.1038/s41557-023-01178-1.

(11) Zhang, B.; Song, W.; Pang, P.; Lai, H.; Chen, Q.; Zhang, P.; Lindsay, S. Role of Contacts in Long-Range Protein Conductance. Proceedings of the National Academy of Sciences 2019, 116 (13), 5886–5891. 10.1073/pnas.1819674116.

(12) Xiang, L.; Zhang, P.; Liu, C.; He, X.; Li, H. B.; Li, Y.; Wang, Z.; Hihath, J.; Kim, S. H.; Beratan, D. N.; Tao, N. Conductance and Configuration of Molecular Gold-Water-Gold Junctions under Electric Fields. Matter 2020, 3 (1), 166–179. 10.1016/j.matt.2020.03.023.

(13) Scullion, L. E.; Leary, E.; Higgins, S. J.; Nichols, R. J. Single-Molecule Conductance Determinations on HS(CH 2) 4 O(CH 2) 4 SH and HS(CH 2) 2 O(CH 2) 2 O(CH 2) 2 SH, and Comparison with Alkanedithiols of the Same Length. Journal of Physics: Condensed Matter 2012, 24 (16), 164211. 10.1088/0953-8984/24/16/164211.

(14) Nishino, T.; Shiigi, H.; Kiguchi, M.; Nagaoka, T. Specific Single-Molecule Detection of Glucose in a Supramolecularly Designed Tunnel Junction. Chemical Communications 2017, 53 (37), 5212–5215. 10.1039/C6CC09932G.

(15) Li, H.; Garner, M. H.; Shangguan, Z.; Chen, Y.; Zheng, Q.; Su, T. A.; Neupane, M.; Liu, T.; Steigerwald, M. L.; Ng, F.; Nuckolls, C.; Xiao, S.; Solomon, G. C.; Venkataraman, L. Large Variations in the Single-Molecule Conductance of Cyclic and Bicyclic Silanes. J Am Chem Soc 2018, 140 (44), 15080–15088. 10.1021/jacs.8b10296.

(16) Chen, H.; Hou, S.; Wu, Q.; Jiang, F.; Zhou, P.; Zhang, L.; Jiao, Y.; Song, B.; Guo, Q.-H.; Chen, X.-Y.; Hong, W.; Lambert, C. J.; Stoddart, J. F. Promotion and Suppression of Single-Molecule Conductance by Quantum Interference in Macrocyclic Circuits. Matter 2021, 4 (11), 3662–3676. 10.1016/j.matt.2021.08.016.

(17) Wang, Z.; Palma, J. L.; Wang, H.; Liu, J.; Zhou, G.; Ajayakumar, M. R.; Feng, X.; Wang, W.; Ulstrup, J.; Kornyshev, A. A.; Li, Y.; Tao, N. Electrochemically Controlled Rectification in Symmetric Single-Molecule Junctions. Proceedings of the National Academy of Sciences 2022, 119 (39). 10.1073/pnas.2122183119.

(18) Wang, Y.; Alangari, M.; Hihath, J.; Das, A. K.; Anantram, M. P. A Machine Learning Approach for Accurate and Real-Time DNA Sequence Identification. BMC Genomics 2021, 22 (1), 525. 10.1186/s12864-021-07841-6.

(19) Komoto, Y.; Ryu, J.; Taniguchi, M. Machine Learning and Analytical Methods for Single-Molecule Conductance Measurements. Chemical Communications 2023, 59 (45), 6796–6810. 10.1039/D3CC01570J.

(20) Chang, S.; Huang, S.; Liu, H.; Zhang, P.; Liang, F.; Akahori, R.; Li, S.; Gyarfas, B.; Shumway, J.; Ashcroft, B.; He, J.; Lindsay, S. Chemical Recognition and Binding Kinetics in a Functionalized Tunnel Junction. Nanotechnology 2012, 23 (23), 235101. 10.1088/0957-4484/23/23/235101.

(21) Fu, T.; Zang, Y.; Zou, Q.; Nuckolls, C.; Venkataraman, L. Using Deep Learning to Identify Molecular Junction Characteristics. Nano Lett 2020, 20 (5), 3320–3325. 10.1021/acs.nanolett.0c00198.

(22) Ryu, J.; Komoto, Y.; Ohshiro, T.; Taniguchi, M. Single-Molecule Classification of Aspartic Acid and Leucine by Molecular Recognition through Hydrogen Bonding and Time-Series Analysis. Chem Asian J 2022, 17 (13). 10.1002/asia.202200179.

(23) Stefani, D.; Guo, C.; Ornago, L.; Cabosart, D.; El Abbassi, M.; Sheves, M.; Cahen, D.; van der Zant, H. S. J.Conformation-Dependent Charge Transport through Short Peptides. Nanoscale 2021, 13 (5), 3002–3009. 10.1039/D0NR08556A.

(24) Tao, S.; Zhang, Q.; Pitie, S.; Liu, C.; Fan, Y.; Zhao, C.; Seydou, M.; Dappe, Y. J.; Nichols, R. J.; Yang, L. Revealing Conductance Variation of Molecular Junctions Based on an Unsupervised Data Analysis Approach. Electrochim Acta 2023, 449, 142225. 10.1016/j.electacta.2023.142225.

(25) Lin, D.; Zhao, Z.; Pan, H.; Li, S.; Wang, Y.; Wang, D.; Sanvito, S.; Hou, S. Using Weakly Supervised Deep Learning to Classify and Segment Single-Molecule Break-Junction Conductance Traces. ChemPhysChem 2021, 22 (20), 2107–2114. 10.1002/cphc.202100414.

(26) Bamberger, N. D.; Ivie, J. A.; Parida, K. N.; McGrath, D. V.; Monti, O. L. A. Unsupervised Segmentation-Based Machine Learning as an Advanced Analysis Tool for Single Molecule Break Junction Data. The Journal of Physical Chemistry C 2020, 124 (33), 18302–18315. 10.1021/acs.jpcc.0c03612.

(27) Aminiranjbar, Z.; Gultakti, C. A.; Alangari, M. N.; Wang, Y.; Demir, B.; Koker, Z.; Das, A. K.; Anantram, M. P.; Oren, E. E.; Hihath, J. Identifying SARS-CoV-2 Variants Using Single-Molecule Conductance Measurements. ACS Sens 2024. 10.1021/acssensors.3c02734.

(28) Tracking SARS-CoV-2 variants. https://www.who.int/en/activities/tracking-SARS-CoV-2-variants/ (accessed 2022-04-06).

(29) Chen, T.; Guestrin, C. XGBoost: A Scalable Tree Boosting System. In Proceedings of the 22nd ACM SIGKDD International Conference on Knowledge Discovery and Data Mining; ACM: New York, NY, USA, 2016; Vol. 42, pp 785–794. 10.1145/2939672.2939785.

(30) Hastie, T.; Friedman, J.; Tibshirani, R. The Elements of Statistical Learning: Data Mining, Inference, and Prediction; Springer Series in Statistics; Springer New York: New York, NY, 2001. 10.1007/978-0-387-21606-5.

(31) Friedman, J. H. Greedy Function Approximation: A Gradient Boosting Machine. The Annals of Statistics 2001, 29 (5), 1189–1232. 10.1214/aos/1013203451.

(32) XGBoost Python Package — xgboost 1.3.0-SNAPSHOT documentation. https://xgboost.readthedocs.io/en/latest/python/index.html (accessed 2020-09-18).

(33) Lecun, Y.; Bottou, L.; Bengio, Y.; Haffner, P. Gradient-Based Learning Applied to Document Recognition. Proceedings of the IEEE 1998, 86 (11), 2278–2324. 10.1109/5.726791.

(34) Kingma, D. P.; Ba, J. Adam: A Method for Stochastic Optimization. 2014.

(35) Wang, Y.; Wang, H.; Das, A. K.; Anantram, M. P. SMBJClassifier. https://github.com/ethanwyr/SMBJClassifier 2024. https://github.com/ethanwyr/SMBJClassifier (accessed 2024-02-28).

