## Supplementary Material for "A ML Framework for Genetic Sequence Identification using 2D Electrical Conductance Probability Distributions from Mixed Data Sets"

### 1. Data acquisition

We relied on SMBJ measurements to capture the conductance characteristics of the DNA sequences. Generally, these experiments (preparing the RNA-DNA hybrids and obtaining conductance traces) take around 4 hours to generate 5,000 conductance traces. We obtained all SMBJ experimental data from [1]. For a detailed explanation and experimental setup, refer to [2], [3]. The experimentally obtained SMBJ current traces are not uniform in duration. This limits the applicability of machine learning classification methods like XGBoost. Therefore, a series of pre-processing steps are used to convert the SMBJ current traces to probability histograms, as explained in the Methods section of the main manuscript and [3]. These histograms have a uniform number of logarithmically spaced bins, introducing uniformity to the current traces. Additionally, a single probability histogram is constructed from multiple traces, thus averaging out the presence of random, high-frequency noise.

### 2. Data visualization

The dataset used contains several thousands of current traces corresponding to ten unique 12 base-pair (bp) short DNA segments from one of three variants of the SARS-CoV-2 genome [1], as detailed in TABLE S.I below. Fig. S1, Fig. S2, Fig. S3 show the empirical *large sample* 1D conductance histograms for our 39 datasets (see TABLE S.I) using an $R^2$ test threshold of $\beta = 0.95$. We refer to these histograms as large sample histograms since these are based on thousands of conductance traces for each dataset. Each row in Fig. S1, Fig. S2, Fig. S3 represents a specific sequence (e.g., Alpha_MM1, Beta_PM, etc.). The variability in the shapes of the histograms shown in each row can be attributed to the inherent stochasticity of the SMBJ experimental procedure and/or variations in experimental parameters such as voltage bias and ramp rate. Similarly, Fig. S4, Fig. S5, Fig. S6 show the empirical large sample 2D conductance histograms for our 39 datasets using an $R^2$ test threshold of $\beta = 0.95$.

TABLE S.I

Summary of the 39 COVID-19 Datasets Used in this Paper

| Label | Sequence | Voltage Bias | Current Amplifier | Ramp Rate |
|---|---|---|---|---|
| E1 | ***Alpha* variant, *mismatch 1*** | 0.10 V | 1 nA/V | 5 V/s |
| E2 | | 0.15 V | 10 nA/V | 3 V/s |
| E3 | 3’- GGG TGA ATA CCA -5’ | 0.10 V | 10 nA/V | 10 V/s |
| E4 | 5’- CCC ACU AAU GGU -3’ | 0.15 V | 10 nA/V | 10 V/s |
| E5 | | 0.20 V | 10 nA/V | 10 V/s |
| E6 | ***Alpha* variant, *mismatch 2*** | 0.15 V | 1 nA/V | 3 V/s |
| E7 | | 0.10 V | 1 nA/V | 10 V/s |
| E8 | 3’- GGG TGA ATA CCA -5’ | 0.10 V | 10 nA/V | 10 V/s |
| E9 | 5’- CCC ACU UAC GGU -3’ | 0.15 V | 10 nA/V | 10 V/s |
| E10 | | 0.20 V | 10 nA/V | 20 V/s |
| E11 | ***Alpha* variant, *perfect m*atch** | 0.05 V | 1 nA/V | 3 V/s |
| E12 | | 0.10 V | 1 nA/V | 3 V/s |
| E13 | 3’- GGG TGA ATA CCA -5’ | 0.10 V | 1 nA/V | 20 V/s |
| E14 | 5’- CCC ACU UAU GGU -3’ | 0.10 V | 10 nA/V | 10 V/s |
| E15 | | 0.20 V | 10 nA/V | 10 V/s |
| E16 | ***Beta variant, mismatch 1*** | 0.10 V | 10 nA/V | 20 V/s |
| E17 | 3’- CCA CAA TTT CCA -5’ | 0.05 V | 10 nA/V | 20 V/s |
| E18 | 5’-GGU GUU GAA GGU-3’ | 0.10 V | 10 nA/V | 20 V/s |
| E19 | ***Beta variant, mismatch 2*** | 0.30 V | 10 nA/V | 20 V/s |
| E20 | 3’- CCA CAA TTT CCA -5’ | 0.30 V | 10 nA/V | 20 V/s |
| E21 | 5’-GGU GUU AAG GGU-3’ | 0.10 V | 10 nA/V | 20 V/s |
| E22 | ***Beta variant, mismatch 3*** | 0.05 V | 10 nA/V | 20 V/s |
| E23 | 3’- CCA CAA TTT CCA -5’ | 0.10 V | 10 nA/V | 20 V/s |
| E24 | 5’-AGU GUU AAA GGU-3’ | 0.10 V | 10 nA/V | 20 V/s |
| E25 | ***Beta variant, perfect match*** | 0.05 V | 1 nA/V | 20 V/s |
| E26 | 3’- CCA CAA TTT CCA -5’ | 0.10 V | 1 nA/V | 20 V/s |
| E27 | 5’-GGU GUU AAA GGU-3’ | 0.10 V | 10 nA/V | 20 V/s |
| E51 | ***Delta variant, mismatch 1*** | 0.03 V | 10 nA/V | 20 V/s |
| E52 | 3’-TCG TTT GGA ACA-5’ | 0.03 V | 10 nA/V | 20 V/s |
| E53 | 5’-AGC ACA CCU UGU-3’ | 0.10 V | 10 nA/V | 20 V/s |
| E54 | | 0.10 V | 10 nA/V | 20 V/s |
| E55 | ***Delta variant, mismatch 2*** | 0.03 V | 10 nA/V | 20 V/s |
| E56 | 3’-TCG TTT GGA ACA-5’ | 0.10 V | 10 nA/V | 20 V/s |
| E57 | 5’-AGC AAG CCU UGU-3’ | 0.10 V | 10 nA/V | 20 V/s |
| E58 | | 0.20 V | 10 nA/V | 20 V/s |
| E59 | ***Delta variant, perfect match*** | 0.03 V | 10 nA/V | 20 V/s |
| E60 | 3’-TCG TTT GGA ACA-5’ | 0.03 V | 10 nA/V | 20 V/s |
| E61 | 5’-AGC AAA CCU UGU-3’ | 0.10 V | 10 nA/V | 20 V/s |
| E62 | | 0.20 V | 10 nA/V | 20 V/s |

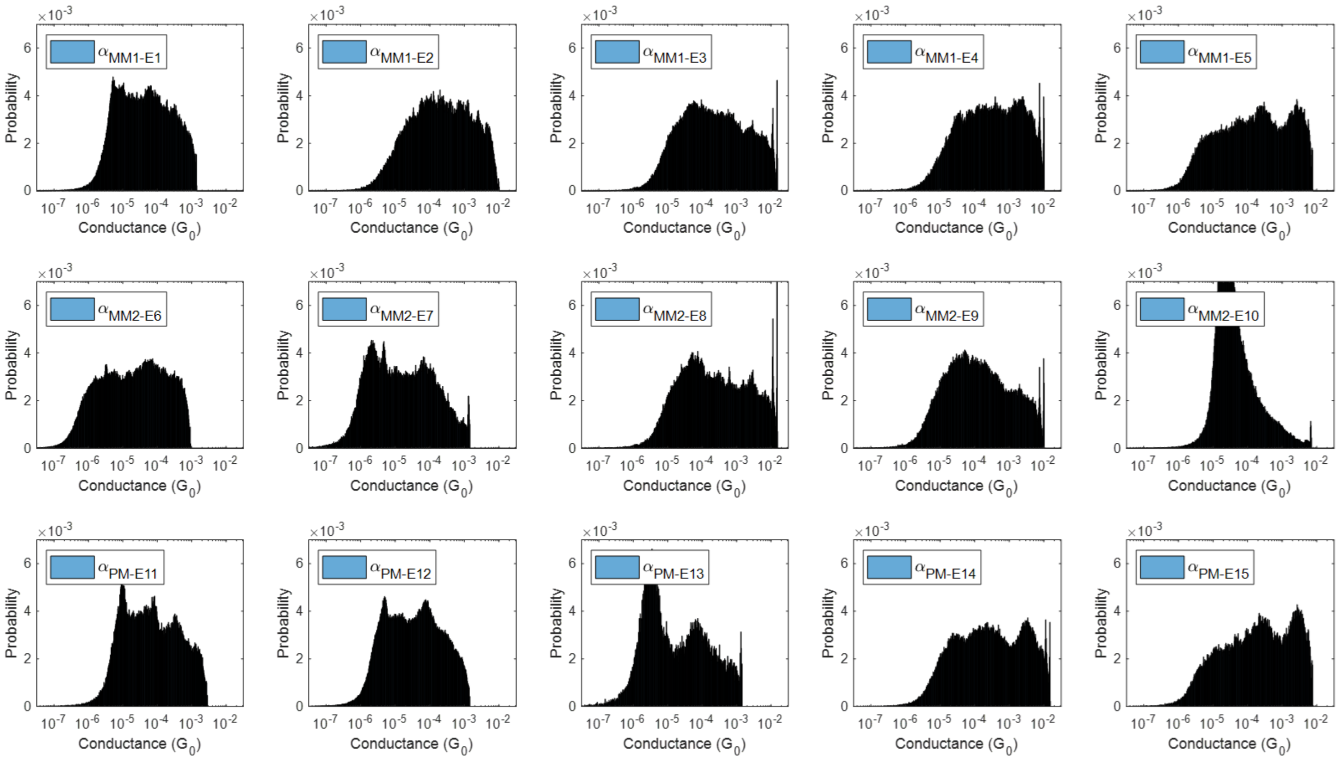


Fig. S1. Empirical large sample 1D conductance histograms for Alpha variant.

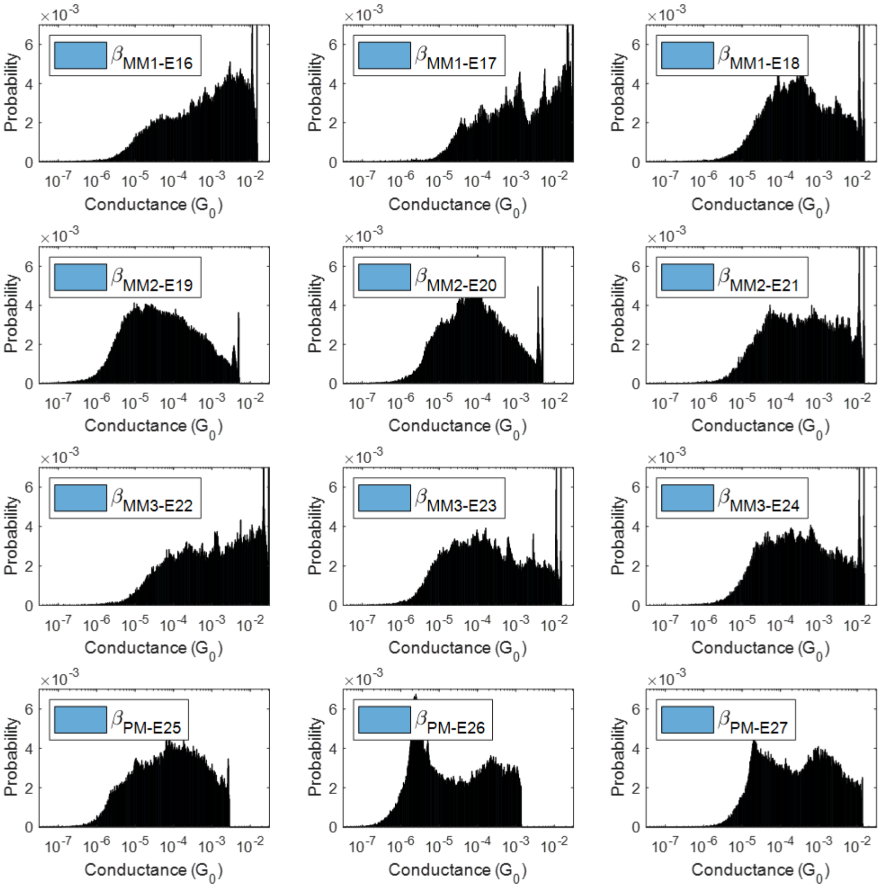


Fig. S2. Empirical large sample 1D conductance histograms for Beta variant.

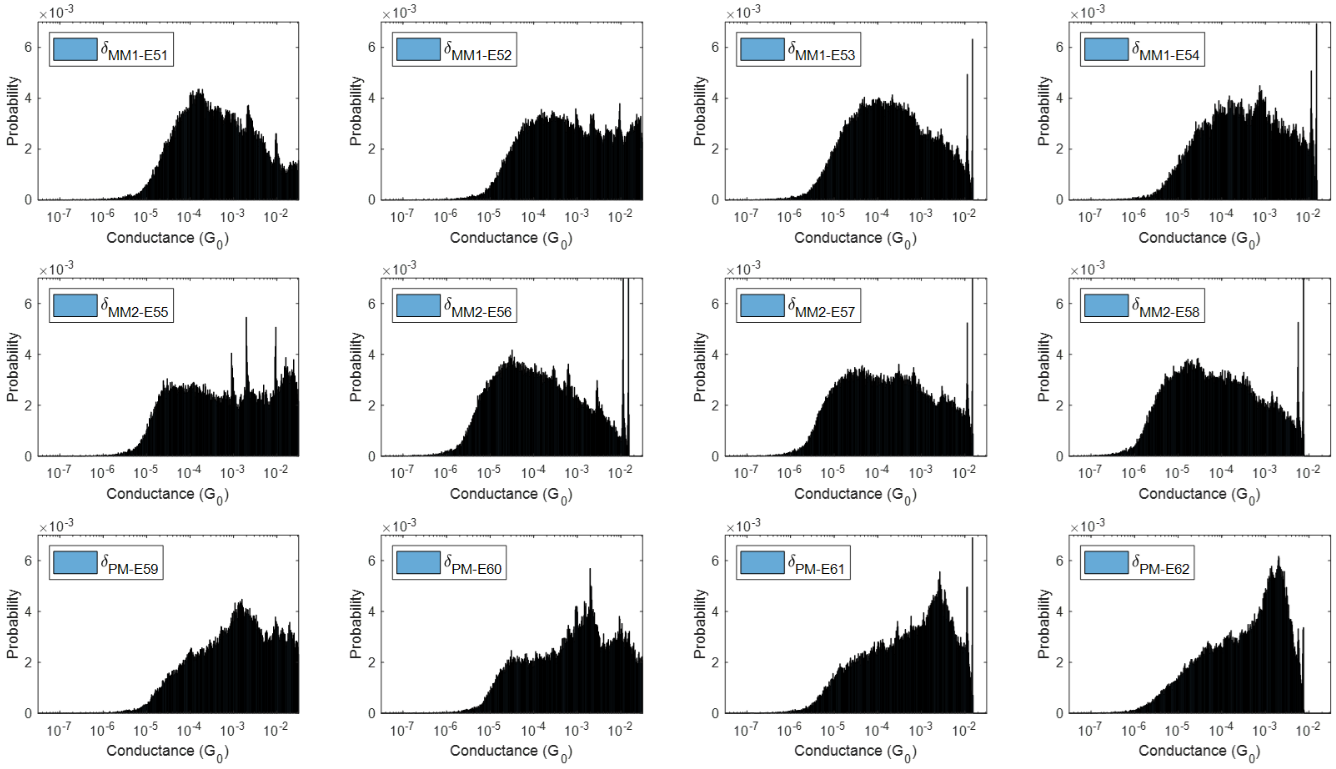


Fig. S3. Empirical large sample 1D conductance histograms for Delta variant.

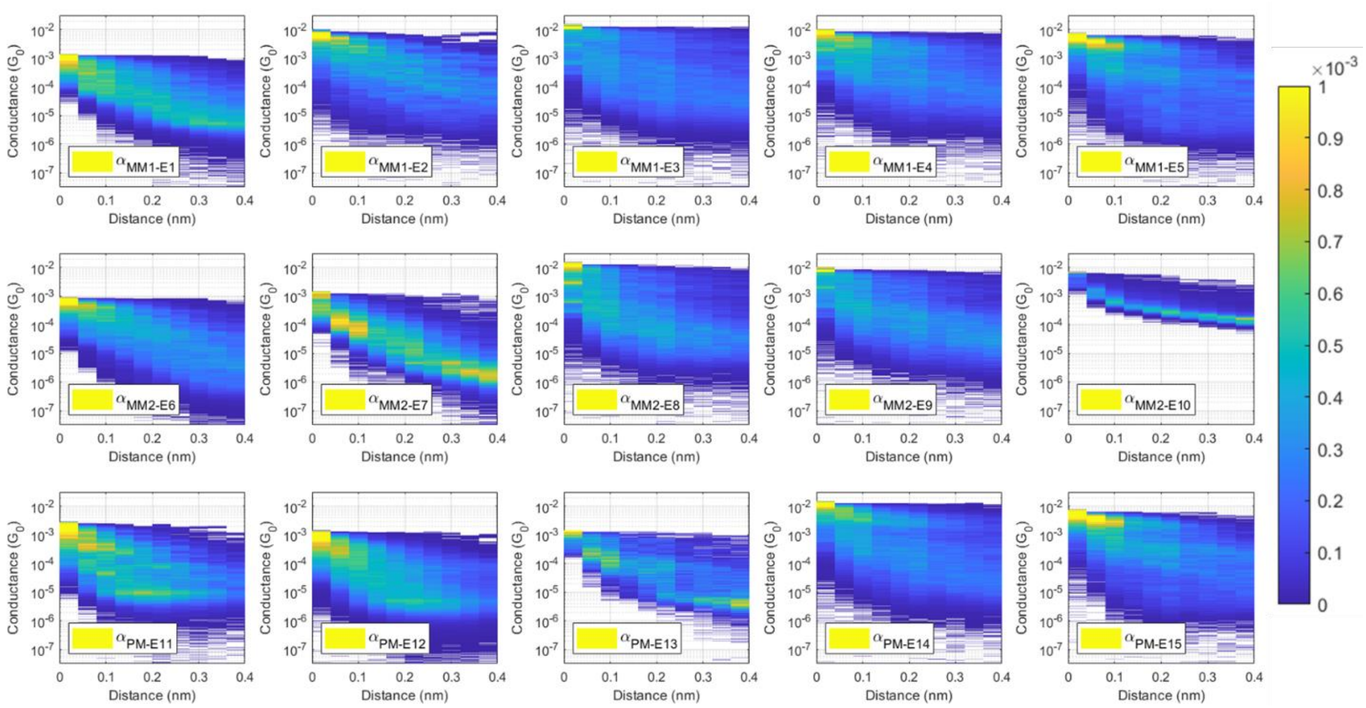


Fig. S4. Empirical large sample 2D conductance histograms for Alpha variant. For better visualization, all elements which have a value larger than 0.001 are represented using the same color scheme as the value 0.001.

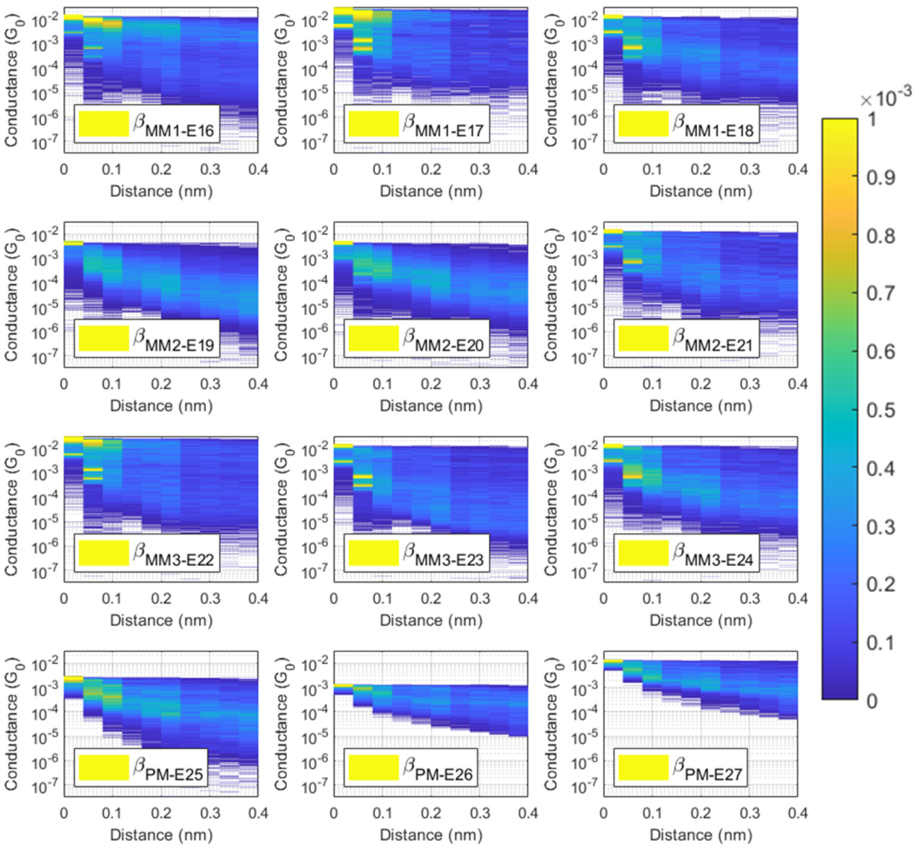


Fig. S5. Empirical large sample 2D conductance histograms for Beta variant. For better visualization, all elements which have a value larger than 0.001 are represented using the same color scheme as the value 0.001.

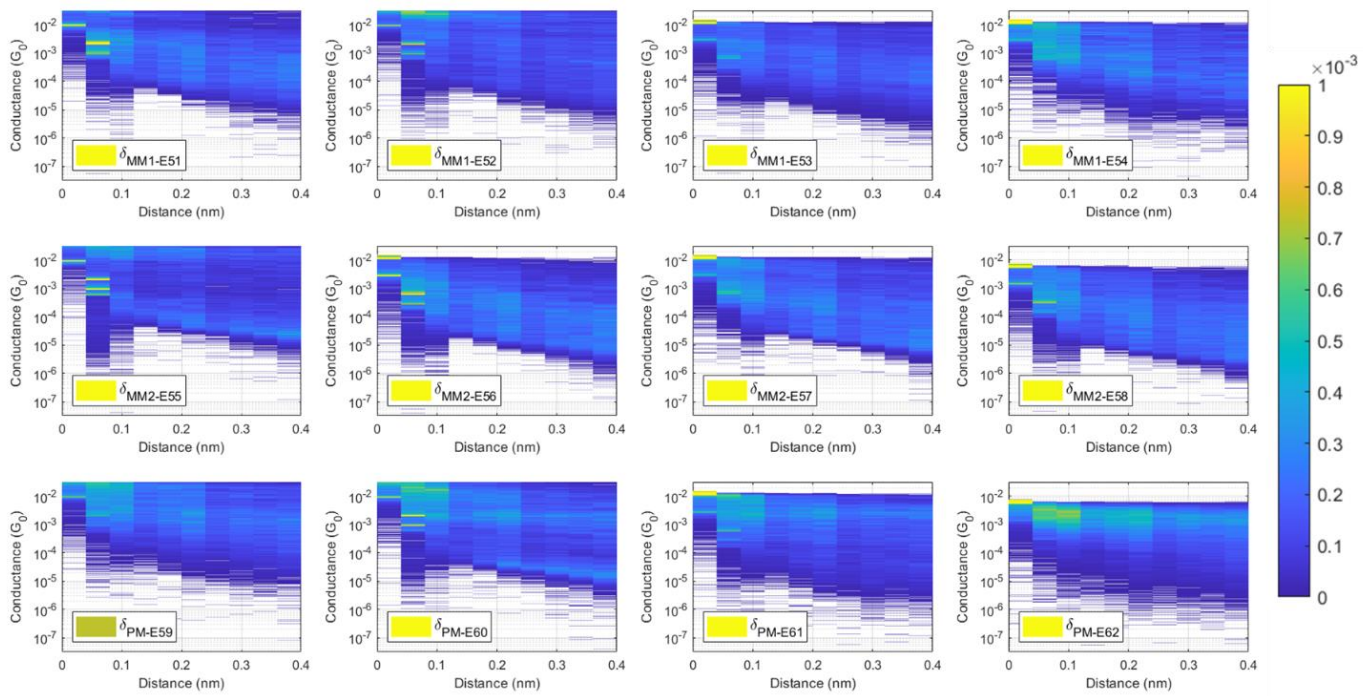


Fig. S6. Empirical large sample 2D conductance histograms for Delta variant. For better visualization, all elements which have a value larger than 0.001 are represented using the same color scheme as the value 0.001.

### 3. Confusion matrices of baseline classifiers

Fig. S7 shows the comparison between the CNN-only approach and the CNN + XGBoost approach. Fig. S8 shows the confusion matrices of baseline classifiers results for all four approaches (A1, A2, A3, A4; see Table I of the main manuscript). The matrices for approaches A1 and A4 have been reproduced in Fig. 1 of the main manuscript.

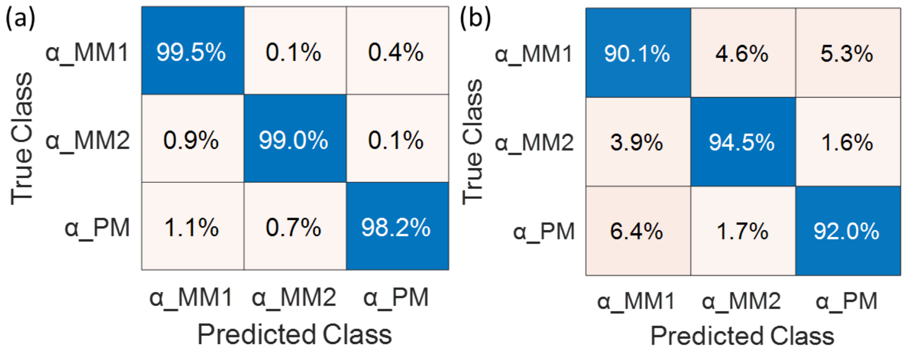


Fig. S7. Confusion matrices for Approach A4 using (a) CNN + XGBoost and (b) CNN only as a machine learning algorithm for classification.

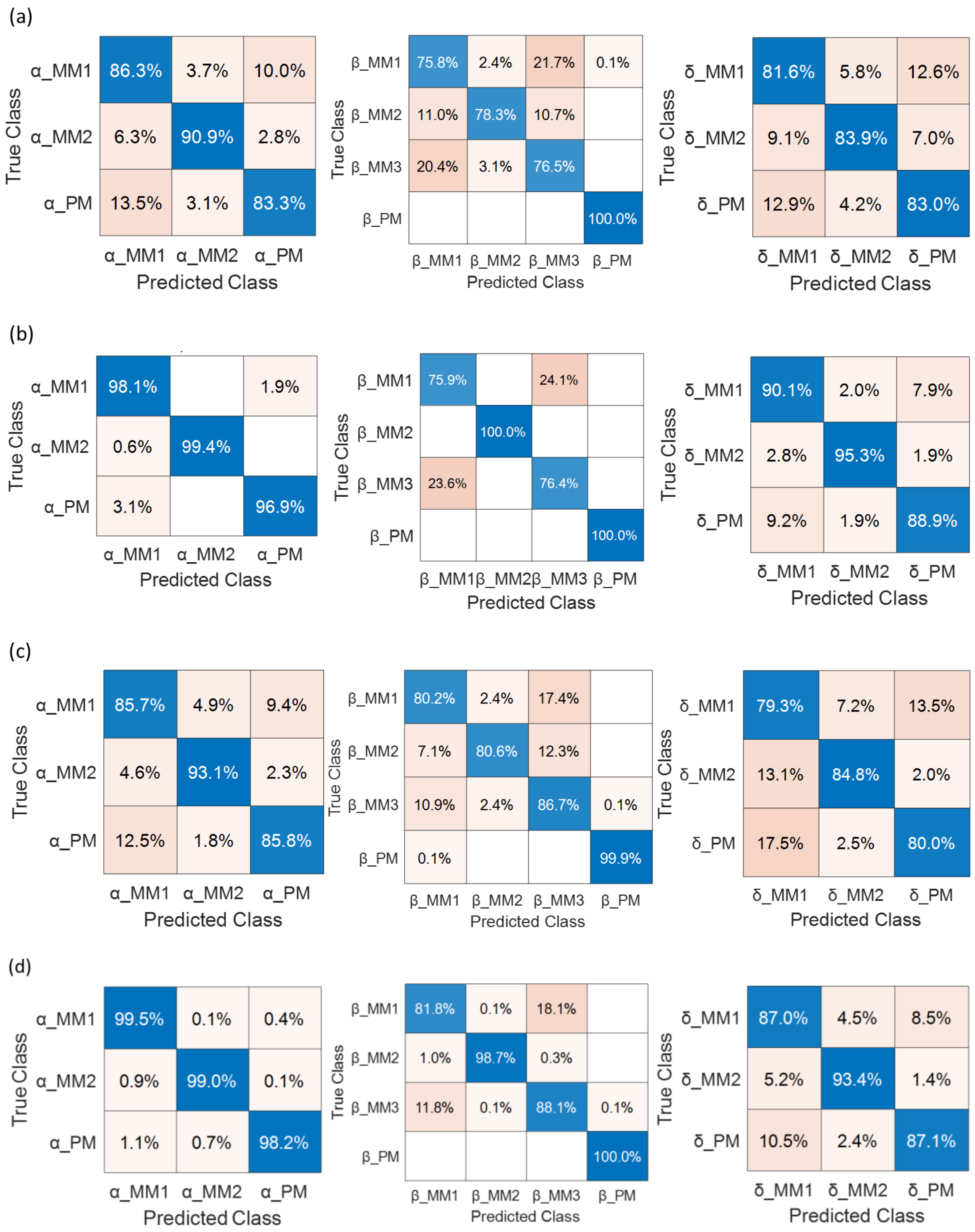


Fig. S8. Confusion matrices for baseline classifiers: (a) Approach A1, (b) Approach A2, (c) Approach A3, and (d) Approach A4. The differences between these four approaches are summarized in Table I of the main manuscript.

Fig. S9 shows the baseline classifier accuracies for the four approaches as a function of $H, H \in [10, 100]$, the number of conductance traces used to construct a histogram. Fig. 2 of the main manuscript shows the same plots but over a smaller range of $H, H \in [10, 50]$.

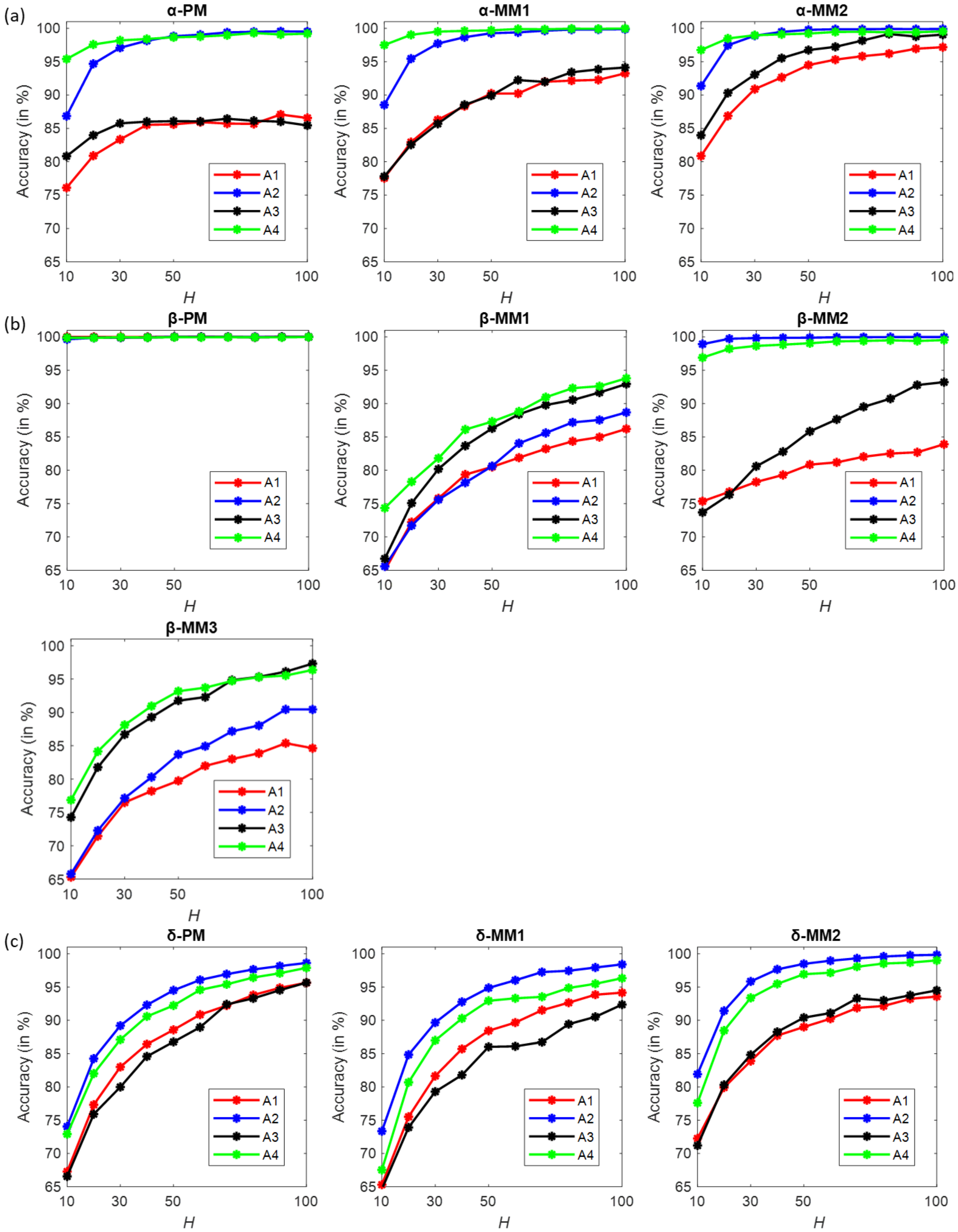


Fig. S9. Performance analysis of baseline classifiers with respect to $H$ for the four approaches.

### 4. Impact of applied bias on classifier accuracy

Fig. S10, Fig. S11, Fig. S12 show the confusion matrices of the Alpha, Beta, and Delta classifiers when the datasets are grouped according to voltage bias. The diagonal elements of these matrices were used to generate Table II of the main manuscript.

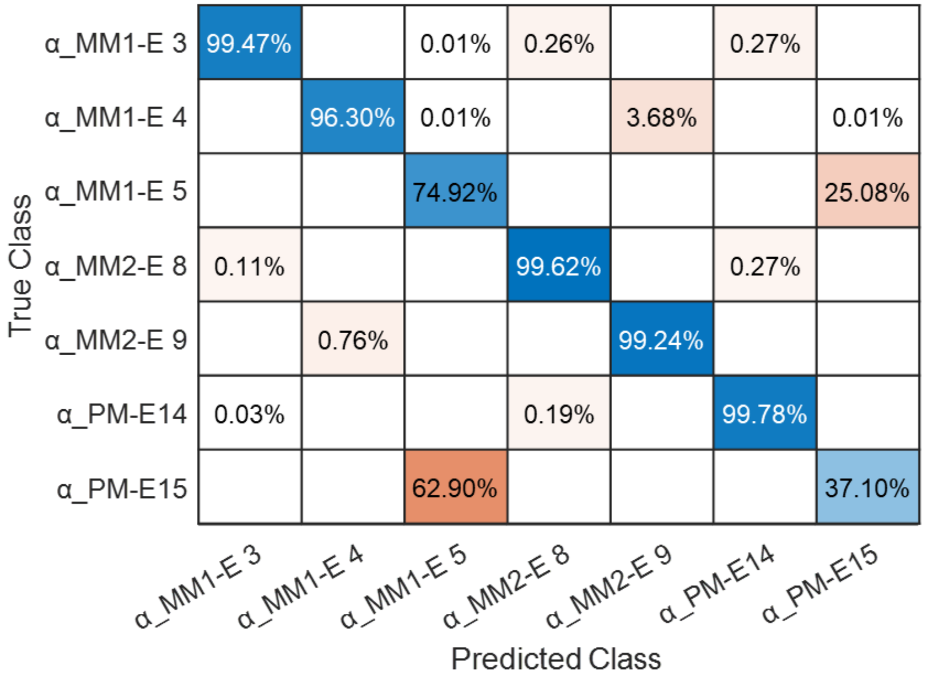


Fig. S10. Confusion matrix of 7-class classifier for Alpha variant (approach A3, $H = 100$).

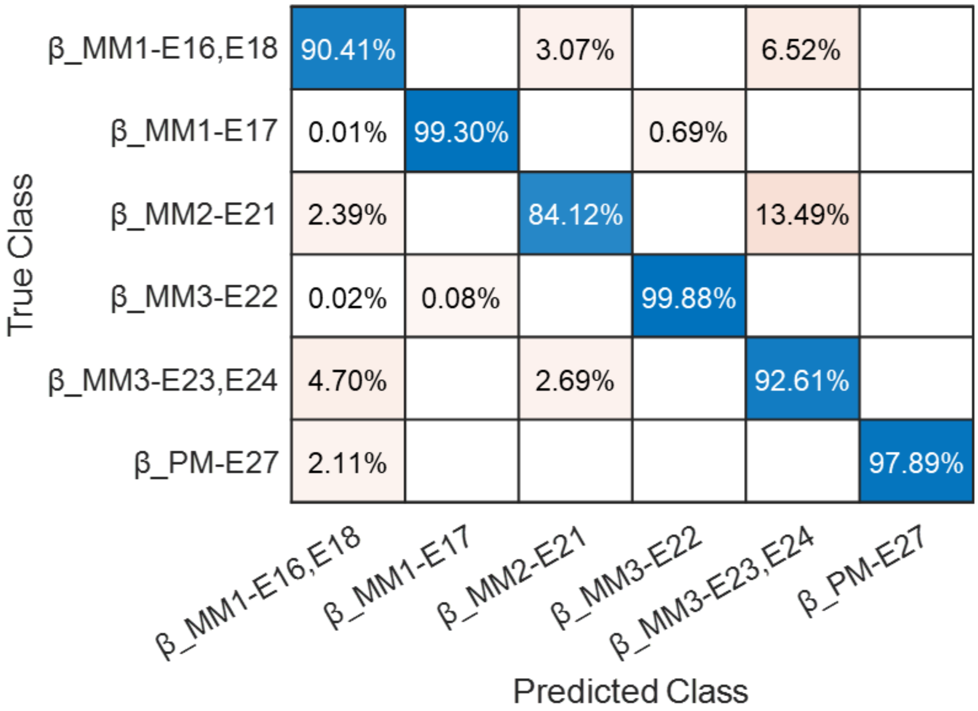


Fig. S11. Confusion matrix of 6-class classifier for Beta variant (approach A3, $H = 100$).

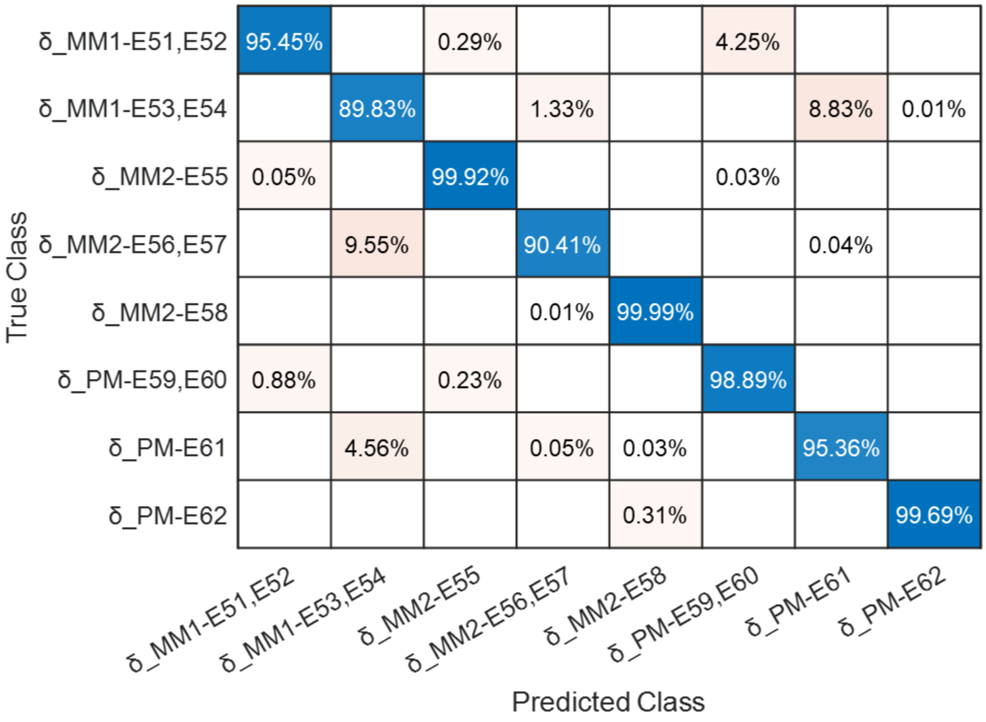


Fig. S12. Confusion matrix of 8-class classifier for Delta variant (approach A3, $H = 100$).

TABLE S.II shows the classifier accuracies for Alpha, Beta, and Delta variants as a function of applied bias using approach A3 and $H = 30$. A similar table for $H = 100$ appears in the main manuscript. The trend observed from Table II of the main manuscript (lower biases result in higher accuracies) holds when classifiers are trained with $H = 30$ histograms, although the impact is somewhat obfuscated by sampling-induced uncertainties.

TABLE S.II

CLASSIFIER ACCURACIES FOR ALPHA, BETA, AND DELTA VARIANTS AS A FUNCTION OF APPLIED BIAS

| | | Dataset | Accuracy | Dataset | Accuracy | Dataset | Accuracy |
|---|---|---|---|---|---|---|---|
| Voltage Bias | | 0.10 V | | 0.15 V | | 0.20 V | |
| Alpha variant | MM1 | E3 | 90.03 % | E4 | 83.49 % | E5 | 62.70 % |
| | MM2 | E8 | 90.93 % | E9 | 89.43 % | | |
| | PM | E14 | 91.80 % | | | E15 | 44.29 % |
| Mean accuracy | | | 90.92 % | | 86.46 % | | 53.50 % |

| | | | | | |
|---|---|---|---|---|---|
| Voltage Bias | | 0.05 V | | 0.10 V | |
| Beta variant | MM1 | E17 | 93.66 % | E16 ∪ E18 | 72.60 % |
| | MM2 | | | E21 | 60.97 % |
| | MM3 | E22 | 97.46 % | E23 ∪ E24 | 76.13 % |
| | PM | | | E27 | 99.99 % |
| Mean accuracy | | | 95.56 % | | 77.42 % |

| Voltage Bias | | 0.03 V | | 0.10 V | | 0.20 V | |
|---|---|---|---|---|---|---|---|
| Delta variant | MM1 | E51 ∪ E52 | 81.65 % | E53 ∪ E54 | 66.27 % | | |
| | MM2 | E55 | 96.61 % | E56 ∪ E57 | 77.91 % | E58 | 99.64 % |
| | PM | E59 ∪ E60 | 88.72 % | E61 | 82.12 % | E62 | 99.44 % |
| Mean accuracy | | | 89.00 % | | 75.43 % | | 99.54 % |

All histograms generated using $H = 30$. All datasets referenced in this table were recorded using a current amplifier of sensitivity 10 nA/V and a ramp rate of 10 V/s.

### 5. Validation by Jensen-Shannon metric

In order to better understand the contradictory behavior exhibited by the Alpha and Delta variants at a bias of 0.20 V (see Table II of the main manuscript), we first observe from the confusion matrix shown in Fig. S10 that Alpha variant E5 (Alpha_MM1) is most often confused with E15 (Alpha_PM). Specifically, the probability that E5 is misclassified as E15 is 0.2508, and the probability that E15 is misclassified as E5 is 0.6290. Clearly, such high misclassification rates should imply a high degree of similarity between the histograms of E5 and E15. In order to verify this conjecture, we computed the pairwise Jensen-Shannon (J-S) distances[4] between the E5 and E15 histograms (normalized as probability distributions) and derived the cumulative distribution function (CDF) of J-S distances. Given two discrete probability distributions $p_x$ and $q_x$ on some random variable $x$, let $m_x$ denote the pointwise mean of $p_x$ and $q_x$, i.e., $m_x = \frac{p_x + q_x}{2}$. Then, the Jensen-Shannon (J-S) distance between $p_x$ and $q_x$ is defined as:

$$\text{J-S distance} = \sqrt{\frac{D(p_x \parallel m_x) + D(q_x \parallel m_x)}{2}},$$

where $D(p_x \parallel m_x)$ denotes the Kullback–Leibler divergence (KLD) of $m_x$ from $p_x$, defined as follows:

$$D(p_x \parallel m_x) = \sum_i p_x(i) \log\left(\frac{p_x(i)}{m_x(i)}\right)$$

While the KLD is not necessarily symmetric, the Jensen-Shannon distance is. In the context of our data, let $p_x$ denote a (1D) conductance distribution for the E5 class and $q_x$ denote a (1D) conductance distribution for the E15 class. Suppose we have 1000 conductance representations for each class. We compute the J-S distances between each pair $\{p_x(m), q_x(n): 1 \leq m, n \leq 1000\}$ and then plot the cumulative distribution function (CDF) of the computed J-S distances. This CDF plot is shown in Fig. S13(a). Since the entire probability mass of J-S distance is concentrated within the band $[0.120, 0.175]$, we can infer that there is substantial similarity between the E5 and E15 histograms, which helps explain the high misclassification rates.

In contrast, we observe from the confusion matrix in Fig. S12 that Delta variant E58 (Delta_MM2) is very occasionally confused with E56 ∪ E57 (Delta_MM2), which occurs with a probability of 0.0001. Similarly, Delta variant E62 (Delta_PM) is very occasionally confused with E58 (Delta_MM2) and occurs with a probability of 0.0031. These extremely small misclassification rates should imply a low degree of similarity between the histograms of (i) E58 and E56 ∪ E57,

(ii) E62 and E58, and (iii) E62 and E61. This conjecture can be verified by examining the corresponding CDF plots of J-S distances shown in Fig. S13(b). We observe that the CDF plots in this case have shifted to the right and the probability mass of J-S distance is almost entirely concentrated within the band $[0.290, 0.485]$. Since the trailing edge of this distance band has shifted to the right compared to what we observe for the Alpha variant, we can infer that the degree of similarity between the "problematic" Delta classes is relatively small, which helps explain the small misclassification rates for E58 and E62.

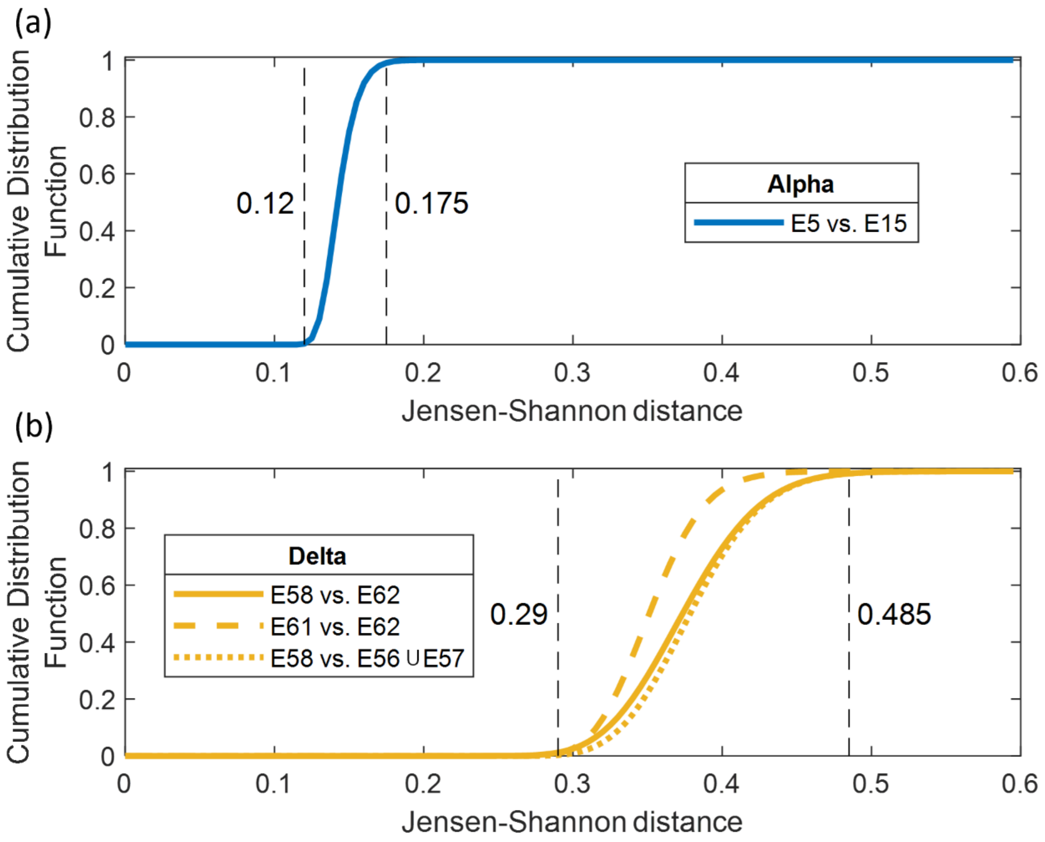


Fig. S13. Cumulative distribution functions (CDF) of Jensen-Shannon distances for (a) Alpha variant datasets E5 and E15, both tested at 0.20 V and (b) Delta variant datasets E58 and E61, both tested at 0.20 V. For the Alpha variant, since E5 is most confused with E15 (see Fig. S10), we show the CDF of the Jensen-Shannon distance between E5 and E15. For the Delta variant, since E58 is very occasionally confused with (E56 ∪ E57) and E62 (see Fig. S12), we show the CDF of the Jensen-Shannon distance between E58 and (E56 ∪ E57), and, between E58 and E62. Additionally, since E61 is very occasionally confused with E62 (see Fig. S12), we show the CDF of the Jensen-Shannon distance between E61 and E62.

**Reference**

[1] Z. Aminiranjbar *et al.*, “Identifying SARS-CoV-2 Variants Using Single-Molecule Conductance Measurements,” *ACS Sens*, May 2024, doi: 10.1021/acssensors.3c02734.

[2] Y. Li *et al.*, “Detection and identification of genetic material via single-molecule conductance,” *Nat Nanotechnol*, vol. 13, no. 12, pp. 1167–1173, Dec. 2018, doi: 10.1038/s41565-018-0285-x.

[3] Y. Wang, M. Alangari, J. Hihath, A. K. Das, and M. P. Anantram, “A machine learning approach for accurate and real-time DNA sequence identification,” *BMC Genomics*, vol. 22, no. 1, p. 525, Dec. 2021, doi: 10.1186/s12864-021-07841-6.

[4] “scipy.spatial.distance.jensenshannon — SciPy v1.12.0 Manual.” Accessed: Jan. 31, 2024. [Online]. Available: https://docs.scipy.org/doc/scipy/reference/generated/scipy.spatial.distance.jensenshannon.html
